# Nanoscopic Clustering of Neuroligin-3 and Neuroligin-4X Regulates Growth Cone Organization and Size

**DOI:** 10.1101/546499

**Authors:** Nicholas J. F. Gatford, P. J. Michael Deans, Rodrigo R.R. Duarte, George Chennell, Pooja Raval, Deepak P. Srivastava

## Abstract

The cell-adhesion proteins neuroligin-3 and neuroligin-4X (NLGN3/4X) have well described roles in synapse formation. NLGN3/4X are also expressed highly during neurodevelopment. However, the role these proteins play during this period is unknown. Here we show that NLGN3/4X localized to the leading edge of growth cones where it promoted neuritogenesis in immature human neurons. Super-resolution microscopy revealed that NLGN3/4X clustering induced growth cone enlargement and influenced actin filament organization. Critically, these morphological effects were not induced by Autism spectrum condition (ASC)-associated NLGN3/4X variants. Finally, actin regulators p21-activated kinase 1 (PAK1) and cofilin were found to be activated by NLGN3/4X and involved in mediating the effects of these adhesion proteins on actin filaments, growth cones, and neuritogenesis. These data reveal a novel role for NLGN3 and NLGN4X in the development of neuronal architecture, which may be altered in the presence of ASD-associated variants.

## Introduction

In mature neurons, axo-dendritic structure is an essential aspect of neuronal function as this is a defining component of neural network connectivity^1^. Development of axo-dendritic morphology begins immediately after neural commitment with the formation and extension of protrusions (neurites) from the cell soma^2^. This process is referred to as neuritogenesis. During this process, immature neurites with actin-rich growth cones form as neural progenitor cells differentiate. Subsequent differentiation leads to the development of the axon and primary dendrite. Extrinsic cues and intrinsic mechanisms work in parallel to orchestrate the development of neuronal morphology^3^. Neuritogenesis is driven by the growth cone, which are highly dynamic and motile subcellular structures^1, 2, 3, 4^. Growth cone behaviour depends on filamentous actin (F-actin) and microtubule dynamics and is regulated by complex signalling pathways^1, 2, 5^. Two major regulators of actin that are also involved in regulating growth cone dynamics are p21-activated kinase (PAK1) and cofilin^2, 6, 7^. PAK1 and cofilin act in concert to enable actin treadmilling and ensure a balance between stable and unstable F-actin in the growth cone. This actin treadmilling provides the growth cone with significant flexibility allowing exploration of the extracellular matrix, response to chemotropic cues, and communication with adjacent cells via cell-adhesion^4^.

Cell-adhesion molecules (CAMs) are also critical regulators of this signalling pathway, operating as molecular clutches between cells which modulate growth cone stabilisation or destabilisation based on cell-cell or cell-matrix interactions^1, 2, 8^. Accumulating evidence suggests that the neurexin-neuroligin cell-adhesion complex (NRXN-NLGN) regulates axonal development^9, 10^,^11^. The NLGNs are a family of post-synaptic proteins which bind to pre-synaptic NRXN; enabling trans-synaptic adhesion, synapse maturation, and establishing synaptic identity in an activity-dependent manner^12, 13, 14^. To date, most studies examining NLGN function have focused on mature neurons, particularly at synapses. Little attention has been given to examining the role of the NLGNs during early neuronal development, despite clear developmental expression of NLGNs^10, 15, 16^. However, there are some key exceptions. For example, NLGN1 and NRXN have been found to contribute to dendritogenesis in Xenopus neurodevelopment^17^. Furthermore, NLGN1 clusters at axonal branch points and filopodia tips during Drosophila cellular neurodevelopment where it acts as a stabilizer, ultimately contributing to axonal arborisation^9^. These studies provide substantial insight into the role of NLGN1 in neurodevelopment but do not examine other NLGNs. Interestingly, mutations in the NRXN-NLGN cell-adhesion complex have frequently been associated with neurodevelopmental disorders, and in particular with autism spectrum conditions (ASC)^13^. ASCs are a heterogenous group of neurodevelopmental disorders of unclear and complex genetic origin^18^. Over 1000 genes are currently associated with ASC; many of which have a neurodevelopmental function, particularly in cellular neurodevelopment^19^. The contribution of disruptions in neurodevelopment as a key component of the pathophysiology of ASCs has been highlighted by recent studies using patient-derived induced pluripotent stem cell models^20, 21^. In the NRXN-NLGN cell-adhesion complex, two key mutations in NLGN3 and NLGN4X^22, 23^, have been well described to impact synapse structure as well as function^24^. However, the role of NLGN3/4X in neurodevelopment and whether the ASC-associated mutations of these proteins impact neurodevelopment and/or the development of neuronal architecture is not known.

In this study, we have identified a novel role for NLGN3/4X during early human neurodevelopment. Using a conditionally immortalised human neural progenitor cellline, we demonstrate that young immature neurons express NLGN3/4X, and that these adhesion proteins form nanoscopic clusters at the leading edge of growth cones. ASC-associated mutant variants of these proteins display an altered nanoscopic distribution. Ectopic NLGN3/4X expression promoted neurite outgrowth, coupled with an enlargement of growth cone size. Using super-resolution imaging, we observed that actin filaments reorganised both with regards to structural organisation as well as bundling, as growth cones enlarged. Interestingly, NLGN3/4X ASC mutants did not exert these effects on neurites or growth cones. We further show that PAK1 was required for both NLGN3/4X-mediated growth cone enlargement as well as nanoscopic clustering of NLGN3/4X. Combined, these results suggest that the cell-adhesion molecules NLGN3/4X regulate growth cone structure and organisation via modulation of actin, ultimately promoting neuritogenesis in human neurodevelopment. Critically, the morphological effects on immature neurons induced by NLGN3 and NLGN4X were not recapitulated by ASC-associated mutants, indicating that abnormal regulation of growth cone dynamics and thus neuritogenesis may contribute to ASC pathophysiology.

## Results

### NLGN3 and NLGN4X are expressed in early human neurodevelopment

The neuroligins (NLGN), and their extracellular binding partner neurexins, are expressed in the mature rodent brain and during rodent and chick embryonic development^15, 16^. However, the temporal expression pattern of NLGN3/4X in human neurodevelopment is unknown. Compiled data from the BrainSpan Atlas of the Developing Human Brain^25^ revealed NLGN3 and NLGN4X are both expressed in the human prenatal brain throughout neurodevelopment and revealed NLGN3 is expressed almost twice as highly as NLGN4X **(Supplemental Figure 1)**. To confirm NLGN3/4X expression in immature human neurons, we assessed expression levels of NLGN3/4X mRNA in undifferentiated human neural progenitor cells (hNPCs) and then immature neurons. NLGN3 exhibited a significant 3.34 fold-change increase in endogenous mRNA expression (**Figure 1A**), while NLGN4X exhibited a 1.467 fold-change increase in endogenous mRNA expression (**Figure 1A)**. Next, using antibodies showing specificity for NLGN3 or NLGN4X (**Supplemental Figure 2 & 3**), we then investigated the expression profile of these adhesion molecules at the protein level. Similar to the mRNA expression, both NLGN3/4X protein levels significantly increased as cells adopted a neuronal fate (**Figure 1B**). This supports the BrainSpan data indicating NLGN3/4X are expressed during the prenatal period, and therefore, may have a functional role during this stage of neurodevelopment.

**Figure 1.**
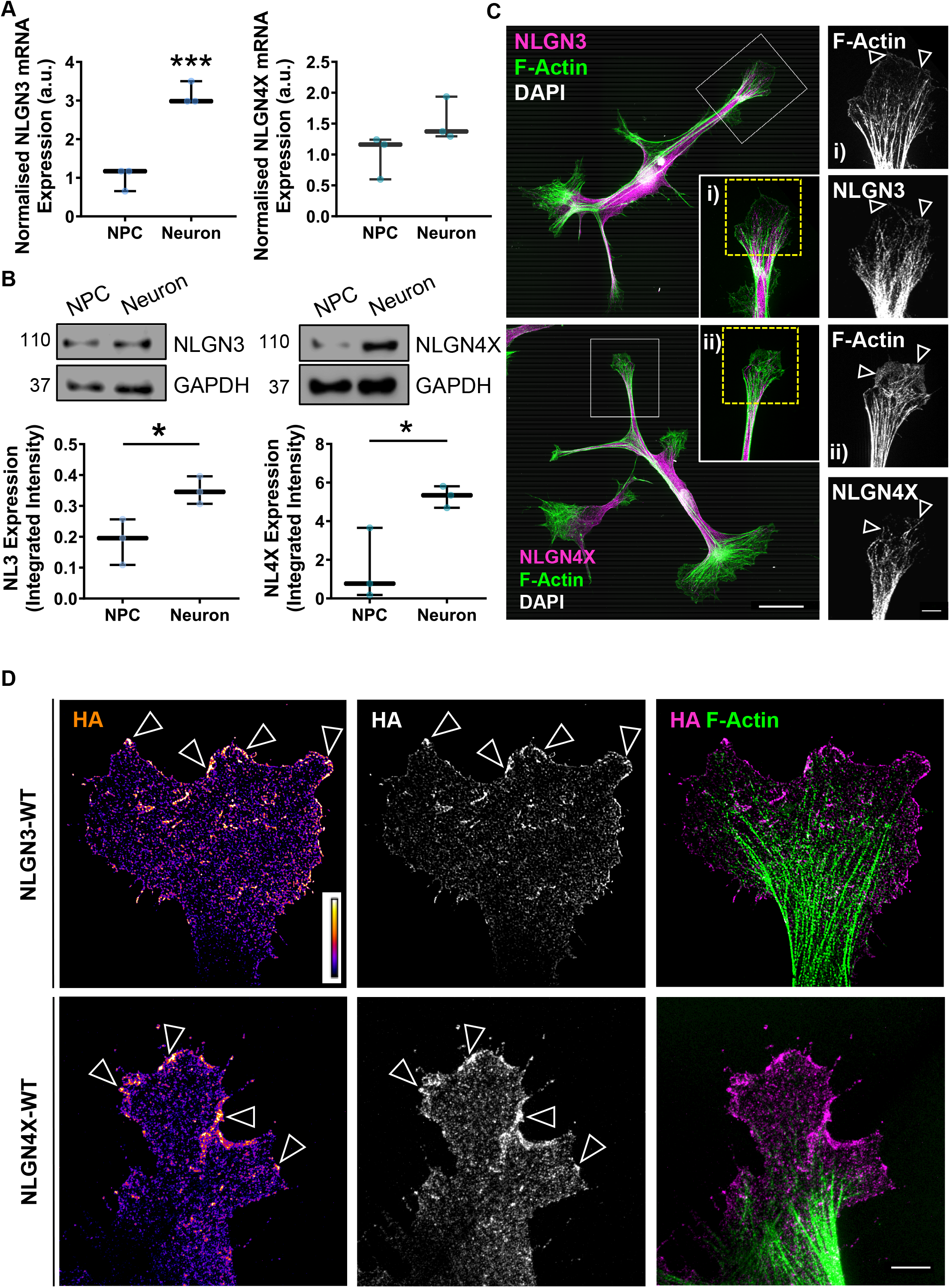
NLGN3 and NLGN4X are expressed in early human neurodevelopment and super-resolution imaging reveals they cluster in nanodomains at the leading edge of growth cones. (A) RT-qPCR data showing endogenous NLGN3 and NLGN4X mRNA expression levels increase as hNPCs differentiate into immature neurons. NLGN3 - t(4)=8.822, p=0.0009, n=3; NLGN4X - t(4)=1.875, p=0.134, n=3. (B) Data and representative western blots showing endogenous NLGN3 and NLGN4X protein expression levels increase as hNPCs differentiate into immature neurons. NLGN3 - t(4)=3.25, p=0.031, n=3; NLGN4X - t(4)=3.35, p=0.029, n=3. (C) Representative super-resolution images of immature human neurons showing endogenous NLGN3 (upper) NLGN4X (lower) expression and localisation. NLGN3 and NLGN4X localise to the growth cone, particularly at the leading edge where they colocalise with F-actin (insets). Scale bar = 25µm (whole cell), 5μm (insets) (D) Representative super-resolution images of human neuronal growth cones ectopically expressing HA-tagged NLGN3 or NLGN4X. Clusters of HA-NLGN3/4X are visible at the growth cone leading edge based on high intensity puncta (open white arrows). Scale bar = 5µm.

### NLGN3 and NLGN4X localize to the leading edge of growth cones

To gain insight into the role of NLGN3/4X in early neurodevelopment, we first assessed the sub-cellular distribution of endogenous NLGN3/4X in immature neurons by super-resolution microscopy. Both NLGN3 and NLGN4X were expressed diffusely throughout the cell. However, discrete puncta for both adhesion proteins could be identified along neurites as well as at the leading edge of growth cones where NLGN3 and NLGN4X colocalized with actin filaments **(Figure 1C, i – ii)**. Previous studies have shown that recruitment and subsequent clustering of NLGNs to synapses is critical for their ability to influence synaptic structure and function^26, 27, 28^. Thus, we reasoned that under the correct conditions, NLGN3/4X would also cluster at the main sites where they exert their effects. To test this, we ectopically expressed HA-tagged wildtype (WT) human NLGN3 or NLGN4X in differentiating neurons for 3 days in order to promote clustering of these adhesion proteins. Both exogenous NLGN3 and NLGN4X were expressed throughout the cell. However, both adhesion proteins were particularly enriched in growth cones, where they could be observed as nanoscopic clusters at the leading edge of growth cones (**Figure 1D)**. Taken together, these data indicate NLGN3 and NLGN4X are expressed at growth cones in immature neurons and upon clustering, enrich at the leading edge of growth cones.

### Mutant NLGN3 and NLGN4X display abnormal sub-cellular distributions

Previous studies have demonstrated that correct subcellular localisation of NLGN3/4X is critical for their ability to influence synaptic structure and function^29, 30, 31^. Interestingly, ASC-associated NLGN3/4X mutants have been reported to display abnormal subcellular distribution compared to wildtype protein in mature neurons^22, 23^. Both mutant proteins, NLGN3-R451C and NLGN4X-D396, are retained intracellularly, and have reduced presence at the cell membrane^22^. To investigate whether this mislocalisation was recapitulated in our cellular system, we first compared the subcellular localisation of ectopic HA-NLGN3/4X and its mutant forms in hNPCs. Whilst HA-NLGN3-WT localised to the plasma membrane, HA-NLGN3-R451C was predominately localised within the cytosol, with only a fraction of the protein at the plasma membrane of hNPCs **(Supplemental Figure 4)**. Similarly, HA-NLGN4X localised to the plasma membrane of hNPCs, whereas HA-NLGN4X-D396 was almost completely localised intracellularly in agreement with previous reports **(Supplemental Figure 4).**

As our data indicated that NLGN3/4X-WT were particularly enriched at growth cones in immature neurons **(Figure 1D),** we next compared the distribution of NLGN3/4X-WT and their mutant variants in immature neurons **(Figure 2A & B)**. Examination of growth cones of immature neurons revealed that NLGN3-R451C was less present at the leading edge but less abundant than the WT form **(Figure 2A)**. Conversely, the NLGN4X-D396 mutant was found to aggregate within the neurite and not localize to the plasma membrane, unlike NLGN4X-WT which localized to the leading edge of growth cones **(Figure 2B)**. Taken together, these data indicate that ASC-associated mutant NLGN3 and NLGN4X display reduced presence at the plasma membranes of hNPCs and abnormal localisation at the leading edge of growth cones in immature neurons.

**Figure 2.**
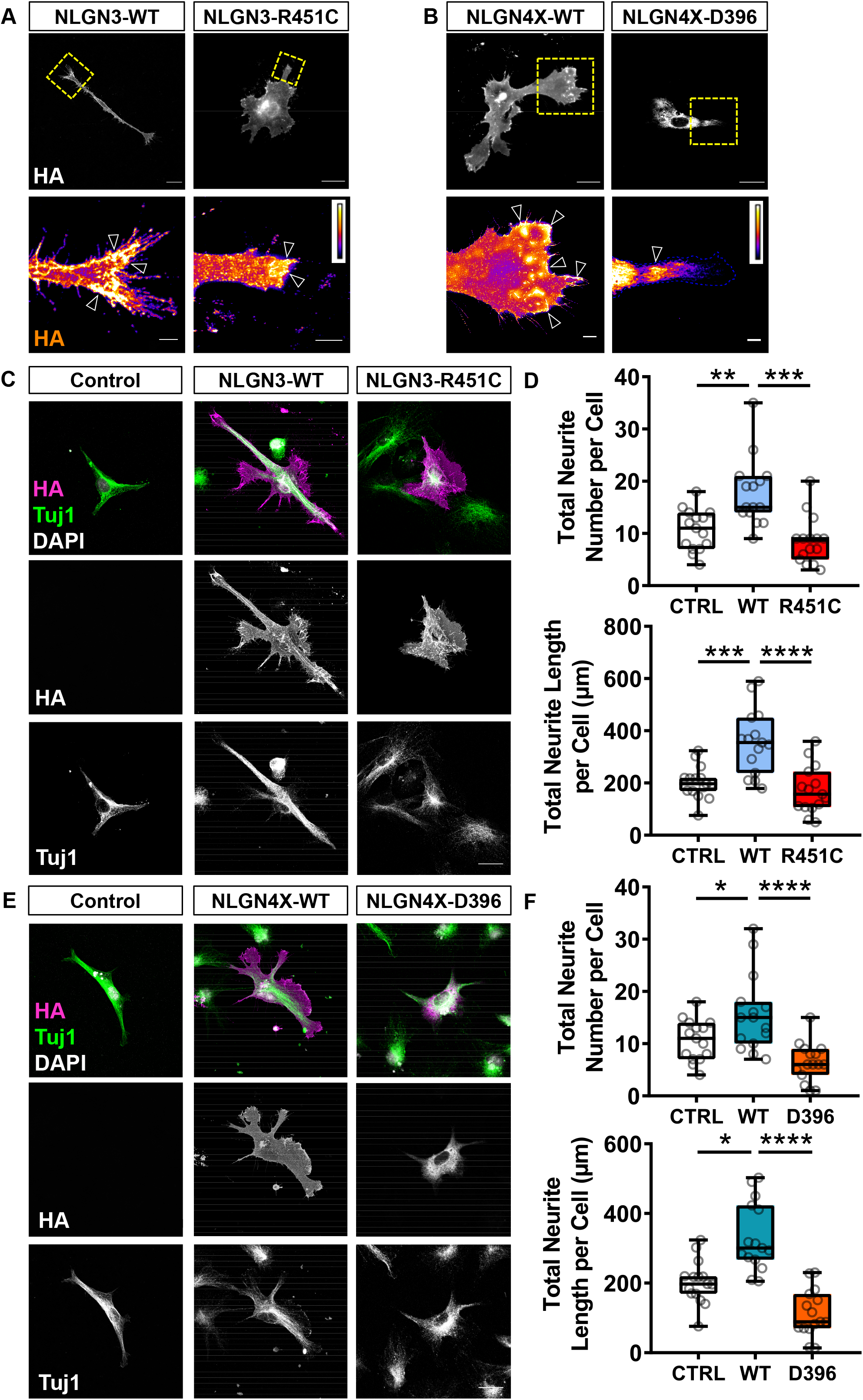
Ectopic wildtype (WT) NLGN3/4X expression in immature neurons increases neuritogenesis in terms of both length and number compared to ectopic ASC-associated NLGN3-R451C or NLGN4X-D396 expression. (A) Representative intensity images showing NLGN3-WT is highly localized to the leading edge of growth cones while the mutant variant is less localized but still present at the leading edge (open white arrows). Scale bar = 5µm (B) Representative intensity images showing NLGN4X-WT is highly localized to the leading edge of growth cones while NLGN4X-D396 is barely present in growth cones. (open white arrows). Scale bar = 5µm (C+D) Representative images and data showing ectopic NLGN3 expression significantly increases neurite number and length in immature neurons. Scale bar = 25µm. Neurite count/cell: Kruskal-Wallis: χ2(3)=18.07, p=0.0001, Dunn: Mean rank difference: −14.37, p=0.008, n=15; neurite length/cell: One-way ANOVA: F(2,42)=15.97, p<0.0001, Bonferroni: t(42)=4.41, p<0.0001, n=15). (E+F) Representative images and data showing ectopic NLGN4X expression significantly increases neurite number and length in immature neurons. Scale bar = 25µm. Neurite count/cell: One-way ANOVA: F(2,42)=11.62, p<0.0001, Bonferroni: t(42)=2.60, p=0.039, n=15; neurite length/cell: Kruskal-Wallis: χ2(3)=26.95, p<0.0001, Dunn: Mean rank difference: −13.53, p=0.014, n=15.

### NLGN3-WT, NLGN4X-WT, and their mutant forms exert differential effects on neurite outgrowth

NLGN3/4X have well established roles in synaptogenesis and spine formation, an effect mediated by the clustering of these proteins at synaptic sites which regulates actin cytoskeletal remodeling^26, 30, 32, 33^. We, therefore, reasoned that as WT NLGN3 and NLGN4X clustered at membranes of hNPCs and at the leading edge of growth cones in immature neurons, that these adhesion proteins may be involved in regulating membrane structures and neurite outgrowth. Indeed, ectopic NLGN3 expression in hNPCs significantly increased cell membrane features not induced by NLGN3-R451C. This could be demonstrated by an increase in the number of lamellipodia on protrusions, which is indicative of cytoskeletal reorganisation at the plasma membrane (count/cell, lamellipodia on protrusions: control, 4.33±0.47; NLGN3-WT, 8.73±2.21; NLGN3-R451C, 6.47±0.76) **(Supplemental Figure 5A & B)**. Next, we examined whether NLGN3-WT or NLGN3-R451C could induce changes in neuritogenesis owing to the enrichment of this adhesion protein at growth cones. Ectopic NLGN3-WT expression significantly increased neuritogenesis in immature neurons (neurite count/cell: control, 10.67±1.01; NLGN3-WT, 17.80±1.67; NLGN3-R451C, 8.60±1.17; neurite length/cell: control, 203.20±16.13 µm; NLGN3-WT, 356.40±31.79 µm; NLGN3-R451C, 173.10±23.36 µm) **(Figure 2C & D)**. This was found to be driven by a significant increase in secondary neuritogenesis i.e. neurites branching from the primary neurite (secondary neurite count/cell: control, 4.07±0.60; NLGN3-WT, 8.20±0.97; NLGN3-R451C, 3.20±0.72); secondary neurite length/cell: control, 58.73±8.61 µm; NLGN3-WT, 152.14±19.78 µm; NLGN3-R451C, 51.99±13.27 µm) **(Supplemental Figure 5C**).

We then investigated whether NLGN4X may also influence cell membrane structures owing to its functional similarity to NLGN3. In hNPCs, we observed that ectopic NLGN4X-WT expression significantly increased cell membrane features not induced by NLGN4X-D396. This could be demonstrated by a decrease in the number of concaves as well as increases in the number of lamellipodia and lamellipodia on protrusions. Conversely, NLGN4X-D396 significantly decreased concaves on cell membranes i.e. sections of inward curving cell membrane (control, 4.87±0.62; NLGN4X-WT, 3.00±0.49; NLGN4X-D396, 1.31±0.29), lamellipodia (control, 2.13±0.32; NLGN4X-D396, 0.39±0.14), and lamellipodia on protrusions (control, 4.33±0.47; NLGN4X-D396, 1.69±0.44) compared to control (**Supplemental Figure 5A & B)**. This suggests the NLGN4X-D396 mutation may operate in a dominant negative way to negatively influence membrane structures.

Next, we ectopically expressed NLGN4X-WT in immature neurons to determine whether this protein also influenced neuritogenesis. Neurons expressing NLGN4X showed a significant increase in neuritogenesis (neurite count/cell: control, 10.67±1.01; NLGN4X-WT, 15.67±1.90; NLGN4X-D396, 6.40±0.96); neurite length/cell: control, 203.20±16.13 µm; NLGN4X-WT, 332.40±25.36 µm; NLGN4X-D396, 114.00±17.40 µm) **(Figure 2E & F)**. This was found to be driven by a significant increase in secondary neuritogenesis (secondary neurite count/cell: control, 4.07±0.60; NLGN4X-WT, 8.20±0.97; NLGN4X-D396, 3.20±0.72); secondary neurite length/cell: control, 58.73±8.61 µm; NLGN4X-WT, 152.14±19.78 µm; NLGN4X-D396, 51.99±13.27 µm) (**Supplemental Figure 5D**). Collectively, these data demonstrate that NLGN3 and NLGN4X can drive morphological changes in hNPCs and significantly promote neurite outgrowth in immature human neurons, consistent with their subcellular distribution. Critically, ASC mutants of these proteins do not exert these effects, which further mirrors their altered subcellular distribution.

### NLGN3 affects growth cone structure by influencing actin filament alignment

The primary driver and navigator of neuritogenesis is the growth cone^4^. The growth cone is a highly dynamic actin-rich structure, therefore, we hypothesized that ectopically expressing NLGN3-WT or NLGN3-R451C would have a striking effect on growth cone morphology and actin filaments. Investigation of growth cones by super-resolution imaging revealed that ectopic NLGN3-WT but not NLGN3-R451C expression significantly increased growth cone area (control, 253.20±28.67 µm^2^; NLGN3-WT, 404.70±51.90 µm^2^; NLGN3-R451C, 218.10±33.31 µm^2^) **(Figure 3A & B)**. This failure of growth cone expansion in the NLGN3-R451C condition may be as a consequence of the decrease in functional NLGN3 clusters at the growth cone leading edge. As growth cone area was found to be significantly correlated to the number of actin filaments within the growth cone **(Supplemental Figure 6A)**, we then looked at whether NLGN3 exerted any influence on actin filaments. Indeed, ectopic NLGN3-WT but not NLGN3-R451C significantly increased growth cone filament number (control, 9.8±1.22; NLGN3-WT, 17.8±1.78; NLGN3-R451C, 8.71±0.60), length, (control, 3.19±0.33 µm; NLGN3-WT, 4.06±0.36 µm; NLGN3-R451C, 3.21±0.18 µm), bundle width (control, 0.36±0.04 µm; NLGN3-WT, 0.84±0.09 µm; NLGN3-R451C, 0.25±0.03 µm), and significantly decreased the distance between growth cone filaments when normalized to filament number (control, 0.16±0.02 µm; NLGN3-WT, 0.06±0.01 µm; NLGN3-R451C, 0.22±0.03 µm) **(Figure 3B)**. Additionally, a significant decrease in anisotropy of actin filaments was found between the NLGN3-WT condition and control (control, 0.16±0.02; NLGN3-WT, 0.07±0.02; NLGN3-R451C, 0.16±0.02); that is, actin filaments in the growth cones of immature neurons ectopically expressing NLGN3-WT are significantly more parallel than actin filaments in immature neurons in control or NLGN3-R451C conditions **(Figure 3A & B)**. Combined, these data suggest NLGN3 has a profound effect on both broad and specific growth cone morphology as well as modulating actin filament orientation within the growth cone.

**Figure 3.**
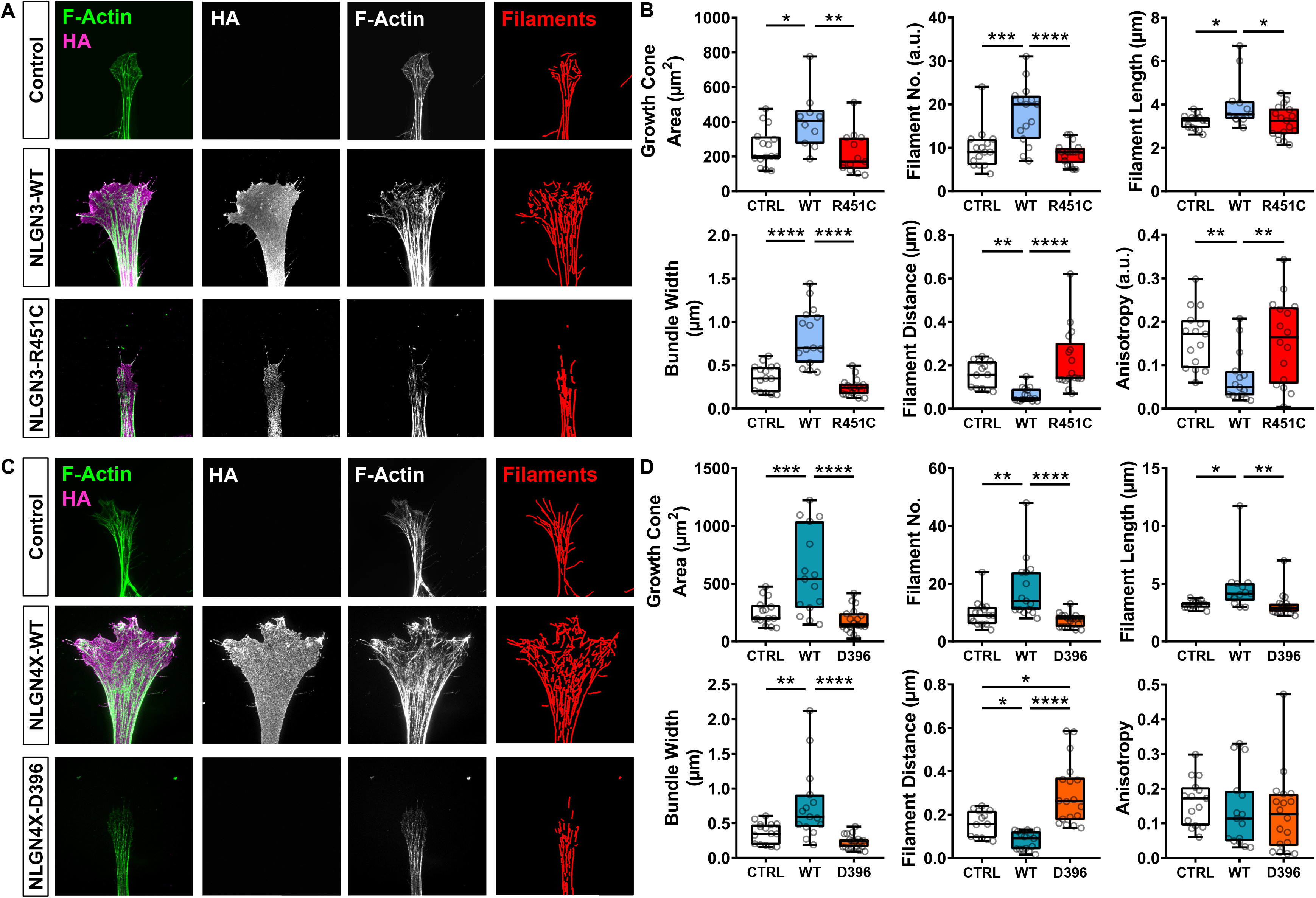
NLGN3 and NLGN4X effects growth cone structure by influencing actin filament organization. (A) Representative super-resolution images showing ectopic wildtype (WT) NLGN3 expression influences growth cone area and actin filament organization. Scale bar = 5µm. (B) Data showing ectopic NLGN3-WT expression significantly increases growth cone area, filament number, filament length, and filament bundle width while decreasing filament distance and anisotropy compared to control or ectopic NLGN3-R451C expression. Growth cone area: One-way ANOVA: F(2,35)=6.434, *p*=0.004, Bonferroni: t(35)=2.86, *p*=0.021, n=15; filament number: One-way ANOVA: F(2,44)=15.57, p<0.0001, Bonferroni: t(44)=4.43, p=0.0002, n=15; filament length: One-way ANOVA: F(2,39)=4.83, p=0.0134, Bonferroni: t(39)=2.73, p=0.028, n=15; bundle width: One-way ANOVA: F(2,44)=33.47, *p*<0.0001, Bonferroni: t(44)=6.10, *p*<0.0001, n=15; filament distance: Kruskal-Wallis: χ2(3)=22.07, p<0.0001, Dunn: Mean rank difference: 17.41, p=0.001, n=15; anisotropy: One-way ANOVA: F(2,43)=6.621, *p*=0.0031, Bonferroni: t(43)=3.21, *p*=0.0075, n=15. (C) Representative super-resolution images showing ectopic NLGN4X-WT expression influences growth cone area and actin filament organization. Scale bar = 5µm. (D) Data showing ectopic NLGN4X-WT expression significantly increases growth cone area, filament number, filament length, and filament bundle width while decreasing filament distance but not anisotropy compared to control or ectopic NLGN4X-D396 expression. Filament distance was also found to be significantly increased between control and ectopic NLGN4X-D396 conditions suggesting a dominant negative effect. Growth cone area: One-way ANOVA: F(2,45)=15.54, p<0.0001, Bonferroni: t(45)=4.22, p=0.0003, n=15; filament number: One-way ANOVA: F(2,46)=12.38, *p*<0.0001, Bonferroni: t(46)=3.51, *p*=0.003, n=15; filament length: Kruskal-Wallis: χ ^2^(3)=16.06, *p*=0.0003, Dunn: Mean rank difference: - 13.47, *p*=0.028, n=15; bundle width: One-way ANOVA: F(2,46)=13.02, *p*<0.0001, Bonferroni: t(46)=3.55, *p*=0.003, n=15; filament distance: Kruskal-Wallis: χ ^2^(3)=27.28, *p*<0.0001, Dunn: Mean rank difference: 12.45, *p*=0.043, n=15, CTRL-NLGN4X-D396 Dunn: Mean rank difference: −12.04, *p*=0.041, n=15; anisotropy: Kruskal-Wallis: χ2(3)=2.59, p=0.2738.

### NLGN4X affects growth cone structure by influencing actin filament organization

We also noticed NLGN4X-WT influencing growth cone morphology and actin filaments in similar ways as NLGN3-WT. We reasoned that these functional similarities are as a consequence of the relatively high protein sequence homology shared between NLGN3 and NLGN4X. Similar to ectopic NLGN3-WT expression, ectopic NLGN4X-WT expression increased growth cone area compared to control (control, 253.20±28.67 µm^2^; NLGN4X-WT, 598.90±95.19 µm^2^; NLGN4X-D396, 182.50±24.85 µm^2^) **(Figure 3C & D)**. No such growth cone expansion was observed in the NLGN4X-D396 condition. This failure of growth cone expansion in the NLGN4X-D396 condition may be as a consequence of the almost total lack of functional NLGN4X clusters at the growth cone leading edge. As growth cone area was found to be significantly correlated to the number of actin filaments within the growth cone **(Supplemental Figure 6B)**, we then looked at whether NLGN4X exerted any influence on actin filaments, similar to NLGN3. Ectopic NLGN4X-WT expression was found to significantly increase growth cone filament number (control, 9.8±1.22; NLGN4X-WT, 18.2±2.74; NLGN4X-D396, 7.21±0.53), length (control, 3.2±0.09 µm; NLGN4X-WT, 4.69±0.62 µm; NLGN4X-D396, 3.14±0.24 µm), bundle width (control, 0.36±0.04 µm; NLGN4X-WT, 0.76±0.14 µm; NLGN4X-D396, 0.23±0.02 µm), and significantly decreased the distance between growth cone filaments when normalized to filament number (control, 0.16±0.02 µm; NLGN4X-WT, 0.08±0.01 µm; NLGN4X-D396, 0.30±0.03 µm) **(Figure 3D)**. Additionally, this data revealed a dominant negative effect for the NLGN4X-D396 mutation as a significant increase in filament distance was found in the NLGN4X-D396 condition compared to control (control, 0.16±0.02 µm; NLGN4X-WT, 0.08±0.01 µm; NLGN4X-D396, 0.30±0.03 µm) **(Figure 3D);** i.e. the NLGN4X-D396 mutation negatively influences the distance between actin filaments within growth cones. However, contrary to the NLGN3-WT anisotropy data, no significant differences in anisotropy were detected between any NLGN4X condition (control, 0.16±0.03; NLGN4X-WT, 0.14±0.03; NLGN4X-D396, 0.13±0.03) **(Figure 3D)**. Taken together, these data suggest NLGN3 and NLGN4X have profound effects on growth cone F-actin remodeling, ultimately leading to significant changes in growth cone size and structure. These data also infer a potential link between NLGN3/4X nanodomain clustering, growth cone F-actin, and actin regulator proteins.

### PAK1 phosphorylation increases in growth cones expressing NLGN3/4X and colocalizes with F-actin

Two major regulators of the actin cytoskeleton that have also been shown to have profound effects on growth cone dynamics/size are p21-activated kinases and the actin severing protein cofilin. Previous studies have shown that PAK1 modulates neurite outgrowth, polarity, and establishing overall neuronal morphology though the regulation of downstream actin regulators within the growth cone^2, 6^. Moreover, cofilin; a further component of the actin treadmilling pathway, has been shown to play a critical role in actin retrograde flow in neurites of the developing brain particularly within growth cones^7^. We therefore hypothesized that the signaling pathway underlying NLGN3 and NLGN4X’s ability to regulate neurite outgrowth and growth cone dynamics may be mediated by PAK1 and/or cofilin. To test this hypothesis, we ectopically expressed NLGN3/4X constructs in HEK293 cells and assessed phospho-PAK1 (pPAK1) and phospho-cofilin (pcofilin) levels, which is indicative of their ability to remodel actin and can therefore be used as proxies for actin dynamics due to their involvement in actin treadmilling^6, 34^. This revealed significant increases in PAK1 and cofilin phosphorylation between the control and NLGN3-WT conditions but no significant difference in phosphorylation for either PAK1/cofilin in the NLGN3-R451C condition **(Figure 4A)**. Significant increases in PAK1 and cofilin phosphorylation were also found between the control and NLGN4X-WT conditions while no significant differences in phosphorylation were found for the NLGN4X-D396 condition **(Figure 4B)**. These changes in PAK1 and cofilin phosphorylation suggest the actin cytoskeletal pathway is activated by NLGN3/4X to induce actin filament reorganization, growth cone enlargement, and ultimately neuritogenesis.

**Figure 4.**
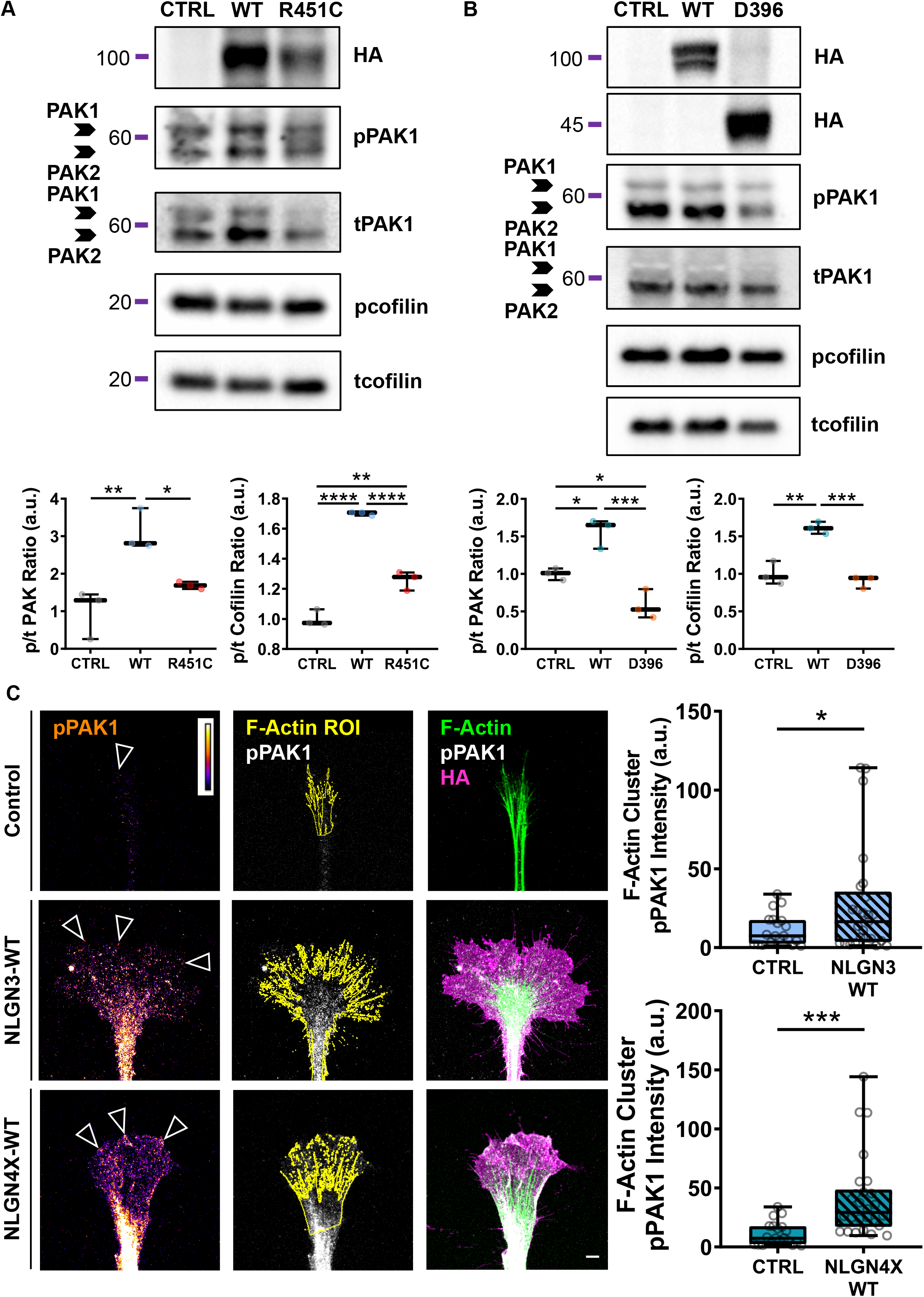
p21-activated kinase (PAK1) phosphorylation increases in growth cones ectopically expressing wildtype (WT) NLGN3 or NLGN4X and colocalizes with growth cone F-actin. (A) Representative blots and data showing ectopic NLGN3-WT expression in HEK293 cells increases phosphorylation of actin regulator proteins PAK1 and cofilin. PAK1: One-way ANOVA: F(2,6)=13.97, p=0.006, Bonferroni: t(6)=5.19, p=0.006, n=3; cofilin: One-way ANOVA: F(2,6)=160.20, *p*<0.0001, Bonferroni: t(6)=17.70, *p*<0.0001, n=3. (B) Representative blots and data showing ectopic NLGN4X-WT expression in HEK293 cells increases phosphorylation of actin regulator proteins PAK1 and cofilin. PAK1: One-way ANOVA: F(2,6)=26.15, p=0.001, Bonferroni: t(6)=4.13, p=0.02, n=3; cofilin: One-way ANOVA: F(2,6)=35.95, *p*=0.0005, Bonferroni: t(6)=6.73, *p*=0.002, n=3. (C) Representative images and data showing phosphorylated PAK1 intensity significantly increases in growth cone F-actin clusters of immature neurons ectopically expressing NLGN3/4X-WT. NLGN3: t(43)=2.07, *p*=0.04 n=25; NLGN4X: t(41)=3.52, *p*=0.001, n=23. Scale bar = 5µm.

To validate that NLGN3/4X increased pPAK1 in immature neurons; thereby indicating a functional interaction, we ectopically expressed NLGN3/4X-WT in immature neurons and measured pPAK1 levels specifically at F-actin/NLGN3/4X clusters. Phospho-PAK1 intensity significantly increased in growth cone F-actin clusters of immature neurons ectopically expressing NLGN3-WT or NLGN4X-WT (NLGN3-WT: control, 11.04±2.22; NLGN3-WT, 27.62±6.92); NLGN4X-WT: control, 11.04±2.22; NLGN4X-WT, 35.39±6.14) **(Figure 4C)**. These data are in line with the immunoblotting data and provide evidence that NLGN3 and NLGN4X strongly induce PAK1 phosphorylation. Taken together, these data suggest PAK1 is involved in the F-actin, growth cone, and neuritogenesis phenotypes demonstrated so far. These data also suggest a mechanistic link between NLGN3/4X, PAK1, and actin dynamics in the growth cone but do not provide causal evidence of a functional relationship.

### Effects of NLGN3/4X on growth cone structure are attenuated by PAK1 inhibition

To demonstrate a causal mechanistic link between NLGN3/4X, PAK1, and growth cone actin dynamics, we ectopically expressed WT NLGN3 or NLGN4X and cotreated with a PAK1 activity inhibitor, FRAX486^35, 36^. Treating immature neurons with 50 nM FRAX486 in the absence of NLGN3/4X significantly decreased growth cone area in control cells compared to vehicle (DMSO) treated cells, as anticipated given the crucial role of PAK1 in growth cone dynamics (**Figure 5A)**. In line with previous data, ectopic NLGN3-WT expression and DMSO treatment resulted in the expected significant increase in growth cone area. Conversely, immature neurons ectopically expressing NLGN3-WT and cotreated with FRAX486, did not display any change in growth cone size compared to control DMSO treated cells (CTRL-DMSO, 172.30±11.02 µm^2^; CTRL-FRAX, 88.02±9.27 µm^2^; NLGN3-DMSO, 468.80±48.98 µm^2^; NLGN3-FRAX, 205.80±19.01 µm^2^) **(Figure 5A & B)**. A similar pattern was observed for filament number i.e. increases in filament number induced by ectopic NLGN3-WT expression are attenuated by FRAX486 cotreatment (CTRL-DMSO, 10.68±0.71; CTRL-FRAX, 3.09±0.33; NLGN3-DMSO, 23.40±1.48; NLGN3-FRAX, 11.95±0.79). Lastly, a similar, although inverse, pattern was observed for decreases in filament distance when normalised to filament number. That is, the decrease in filament distance induced by NLGN3-WT ectopic expression was attenuated by FRAX486 cotreatment (CTRL-DMSO, 0.10±0.01 µm; CTRL-FRAX, 0.44±0.06 µm; NLGN3-DMSO, 0.04±0.004 µm; NLGN3-FRAX, 0.08±0.008 µm) **(Figure 5A & B)**. Taken together, these data strongly implicate PAK1 pathway activation as a core component of the molecular mechanism underlying the previously observed subcellular phenotypes induced by ectopic NLGN3-WT expression and demonstrate a causal association between NLGN3, PAK1, and growth cone actin dynamics.

**Figure 5.**
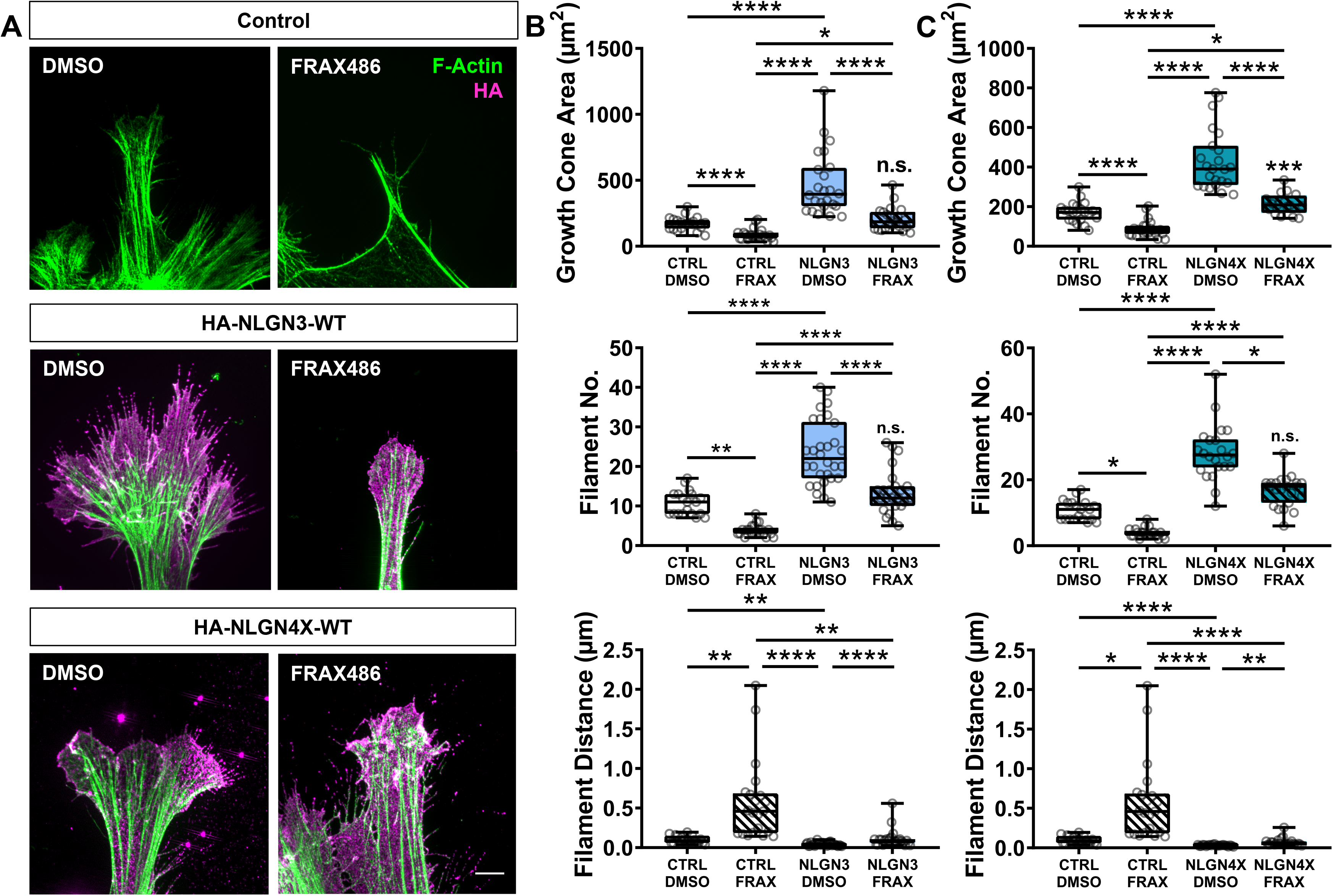
Growth cone morphology changes induced by ectopic NLGN3/4X-WT expression are attenuated by PAK1 inhibition. (A) Representative images showing increases in growth cone area, filament number, and decreases in filament distance induced by ectopic NLGN3/4X-WT expression are attenuated by PAK1 inhibition. Scale bar = 5µm. (B) Data showing increases in growth cone area, filament number, and decreases in filament distance induced by ectopic NLGN3-WT expression are attenuated by PAK1 inhibition. Growth cone area: Kruskal-Wallis: χ^2^(4)=62.71, *p*<0.0001; CTRL-DMSO/CTRL-FRAX - Dunn: Mean rank difference: 24.68, p=0.01, n=25; CTRL-DMSO/NLGN3-DMSO - Dunn: Mean rank difference: −35.21, p<0.0001, n=25; CTRL-DMSO/NLGN3-FRAX - Dunn: Mean rank difference: −6.50, p>0.99, n=25. Filament number: Kruskal-Wallis: χ2(4)=70.82, p<0.0001; CTRL-DMSO/CTRL-FRAX - Dunn: Mean rank difference: 29.17, p=0.003, n=25; CTRL-DMSO/NLGN3-DMSO - Dunn: Mean rank difference: −34.10, p<0.0001, n=25; CTRL-DMSO/NLGN3-FRAX - Dunn: Mean rank difference: −5.33, p>0.99, n=25. Filament distance: Kruskal-Wallis: χ2(4)=60.25, p<0.0001; CTRL-DMSO/CTRL-FRAX - Dunn: Mean rank difference: - 29.84, p=0.003, n=25; CTRL-DMSO/NLGN3-DMSO - Dunn: Mean rank difference: 29.82, p=0.0007, n=25; CTRL-DMSO/NLGN3-FRAX - Dunn: Mean rank difference: 5.75, p>0.99, n=25. (C) Data showing increases in growth cone area, filament number, and decreases in filament distance induced by ectopic NLGN4X-WT expression are attenuated by PAK1 inhibition. Growth cone area: Kruskal-Wallis: χ2(4)=64.28, p<0.0001; CTRL-DMSO/CTRL-FRAX - Dunn: Mean rank difference: 20.78, p=0.03, n=25; CTRL-DMSO/NLGN4X-DMSO - Dunn: Mean rank difference: −35.17, p<0.0001, n=25; CTRL-DMSO/NLGN4X-FRAX - Dunn: Mean rank difference: −10.84, p>0.99, n=25. Filament number: Kruskal-Wallis: χ2(4)=69.94, p<0.0001; CTRL-DMSO/CTRL- FRAX - Dunn: Mean rank difference: 23.17, p=0.02, n=25; CTRL-DMSO/NLGN4X- DMSO - Dunn: Mean rank difference: −37.11, p<0.0001, n=25; CTRL-DMSO/NLGN4X-FRAX - Dunn: Mean rank difference: −15.55, p>0.99, n=25. Filament distance: Kruskal-Wallis: χ2(4)=65.93, p<0.0001; CTRL-DMSO/CTRL-FRAX - Dunn: Mean rank difference: −22.16, p=0.03, n=25; CTRL-DMSO/NLGN4X-DMSO - Dunn: Mean rank difference: 36.72, p<0.0001, n=25; CTRL-DMSO/NLGN4X-FRAX - Dunn: Mean rank difference: 13.90, p>0.99, n=25.

Inhibiting PAK1 has a similar effect on NLGN4X-induced remodeling of growth cones and actin. Consistent with previous data, ectopic NLGN4X-WT expression significantly increased growth cone area. However, cotreatment with FRAX486 (50 nM) attenuated NLGN4X-WT-induced enlargement of growth cone area (CTRL-DMSO, 172.30±11.02 µm^2^; CTRL-FRAX, 88.02±9.27 µm^2^; NLGN4X-DMSO, 432.00±32.09µm^2^; NLGN4X-FRAX, 214.70±13.40 µm^2^) **(Figure 5A & C)**. Cotreatment with FRAX486 also blocked increases in filament number induced by ectopic NLGN4X-WT expression (CTRL-DMSO, 10.68±0.71; CTRL-FRAX, 3.09±0.33; NLGN4X-DMSO, 28.18±1.81; NLGN4X-FRAX, 16.18±0.85). Lastly, the decrease in filament distance induced by ectopic NLGN4X-WT expression is attenuated by FRAX486 cotreatment (CTRL-DMSO, 0.10±0.01 µm; CTRL-FRAX, 0.44±0.06 µm; NLGN4X-DMSO, 0.03±0.002 µm; NLGN4X-FRAX, 0.06±0.006 µm) **(Figure 5A & C)**. Combined, these data implicate PAK1 pathway activation as a core component of the molecular mechanism underlying the previously observed subcellular phenotypes induced by ectopic NLGN4X-WT expression and further demonstrate a causal link between NLGN4X, PAK1, and growth cone actin dynamics.

### PAK1 Inhibition Decreases NLGN3/4X Growth Cone Clustering

We previously observed that wildtype NLGN3/4X formed nanoscopic clusters at the growth cone leading edge where it colocalized with actin. Conversely, mutant forms of these proteins displayed reduced localization in growth cones and impaired ability to remodel growth cone morphology and actin dynamics. Interestingly, inhibiting PAK1 signaling with FRAX486 produced growth cone morphologies and actin dynamics similar to that produced by mutant NLGN3/4X. Therefore, we hypothesised that treatment with FRAX486 may also disrupt the nanoscopic clustering of ectopic NLGN3/4X-WT within growth cones. To examine this, we compared the abundance of either adhesion protein at the leading edge of growth cones with or without FRAX486 cotreatment. This revealed significant decreases in NLGN3-WT growth cone cluster number, area, and intensity in FRAX486 compared to DMSO treated cells (number: NLGN3-DMSO, 863.80±127.50; NLGN3-FRAX, 158.70±15.29); area: NLGN3-DMSO, 0.17±0.04 µm^2^; NLGN3-FRAX, 0.05±0.008 µm^2^); intensity: NLGN3-DMSO, 11.46±3.53; NLGN3-FRAX, 4.38±1.09) **(Figure 6A)**. A similarly significant decrease was also observed for NLGN4X-WT growth cone clusters in terms of number, area, and intensity when treated with FRAX486 compared to DMSO (number: NLGN4X-DMSO, 713.70±83.84; NLGN4X-FRAX, 212.80±35.68); area: NLGN4X-DMSO, 0.21±0.05 µm^2^); NLGN4X-FRAX, 0.06±0.01 µm^2^; intensity: NLGN4X-DMSO, 10.44±2.32; NLGN4X-FRAX, 3.81±0.86) **(Figure 6B)**. Combined, these data suggest PAK1 activation also plays a role in clustering NLGN3/4X at the growth cone promoting its adhesive function. Furthermore, combined with previous data, this not only suggests NLGN3/4X are capable of activating PAK1 in the growth cone but also PAK1 activation can promote NLGN3/4X clustering in growth cones.

**Figure 6.**
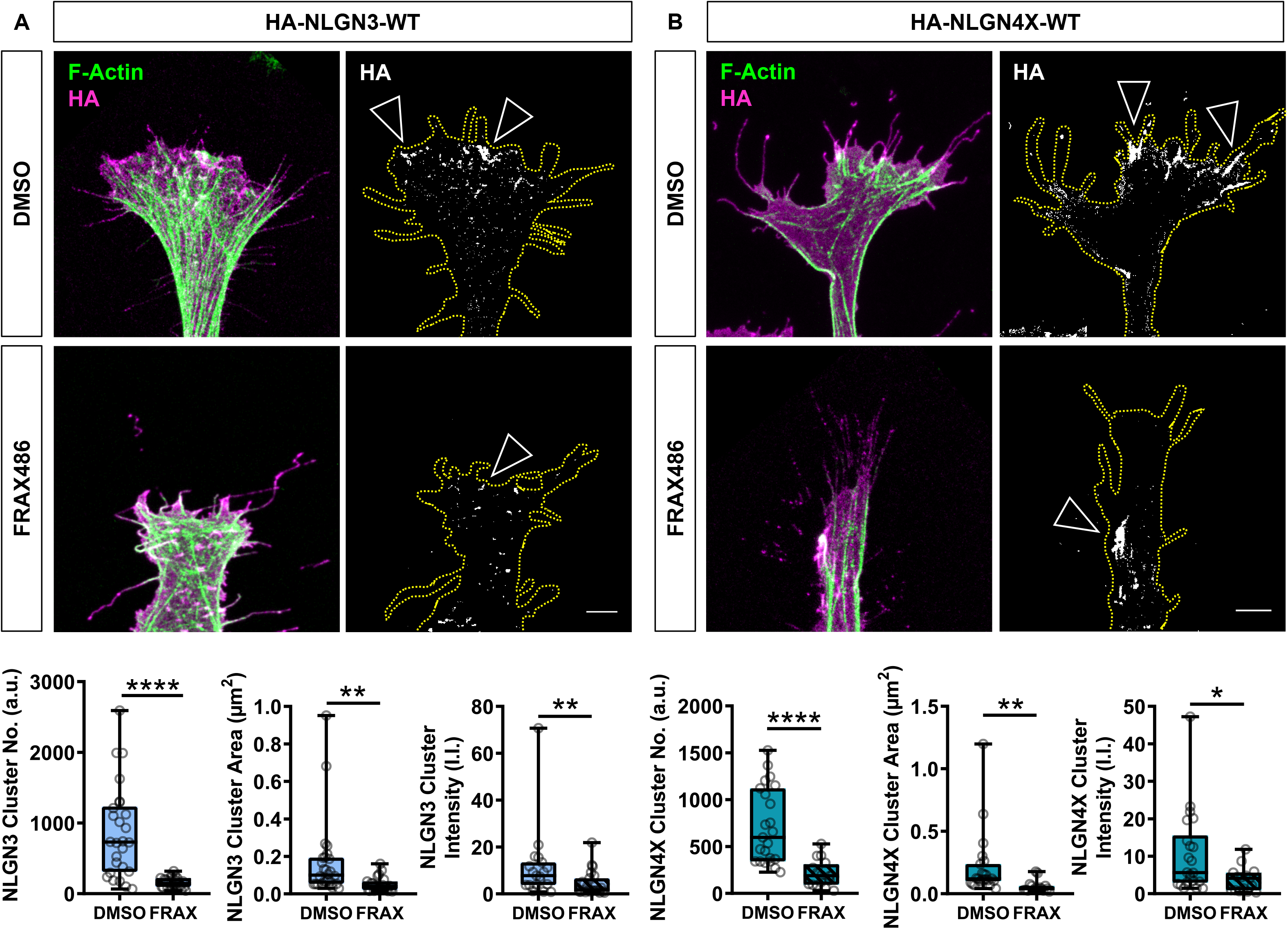
Growth cone NLGN3/4X clusters induced by ectopic HA-NLGN3/4X expression are also attenuated by PAK1 inhibition, a NLGN-PAK1 feedback loop is likely to play a role. (A) Representative images and data showing ectopically expressed HA-NLGN3-WT clusters are attenuated by PAK1 inhibition in terms of cluster number, area, and intensity. Scale bar = 5µm. Number: T-test: t(47)=5.16, p<0.0001, n=25; area: Mann-Whitney: (U=100, p<0.0001, n=25); integrated intensity (I.I.): Mann-Whitney: (U=117, *p*=0.0096, n=25). (B) Representative images and data showing ectopically expressed HA-NLGN4X-WT clusters are attenuated by PAK1 inhibition in terms of cluster number, area, and intensity. Scale bar = 5µm Number: T-test: t(37)=4.76, *p*<0.0001, n=20; area: Mann-Whitney: (U=41, p<0.0001, n=20); integrated intensity (I.I.): Mann-Whitney: (U=87, *p*=0.015, n=20).

### Effects of NLGN3/4X on neurite outgrowth are attenuated by PAK1 inhibition

Given that FRAX486 attenuated the nanoscopic clustering of ectopic NLGN3/4X-WT in growth cones, we hypothesized that this would also impact the overall morphology of the cell. Consistent with this notion, FRAX486 treatment significantly decreased neurite number and length in immature neurons compared to DMSO controls. Similar to previous data, ectopic NLGN3-WT expression resulted in the expected significant increase in neurite outgrowth. However, FRAX486 cotreatment inhibited NLGN3-WT-induced effects on neurite number and length (number: CTRL-DMSO, 7.05±0.43; CTRL-FRAX, 3.14±0.24; NLGN3-DMSO, 14.00±1.16; NLGN3-FRAX, 6.26±0.37; length: CTRL-DMSO, 181.00±8.32 µm; CTRL-FRAX, 59.89±6.00 µm; NLGN3-DMSO, 302.20±22.45 µm; NLGN3-FRAX, 130.40±8.87 µm) **(Figure 7A)**. A similar pattern was observed when the same experimental paradigm was applied to NLGN4X. In agreement with previous data, ectopic NLGN4X-WT expression resulted in significantly increased neurite number and length. However, again, cotreatment with FRAX486 blocked NLGN4X-WT’s effect on neuritogenesis (number: CTRL-DMSO, 7.05±0.43; CTRL-FRAX, 3.14±0.24; NLGN4X-DMSO, 13.61±1.03; NLGN4X-FRAX, 7.31±0.58; length: CTRL-DMSO, 181.00±8.32 µm; CTRL-FRAX, 59.89±6.00 µm; NLGN4X-DMSO, 276.40±15.08 µm; NLGN4X-FRAX, 153.30±13.44 µm) **(Figure 7B)**. Taken together, these data indicate that clustering of NLGN3/4X at the leading edge of growth cones is required for both increased growth cone dynamics and neuritogenesis via a PAK1-dependent mechanism.

**Figure 7.**
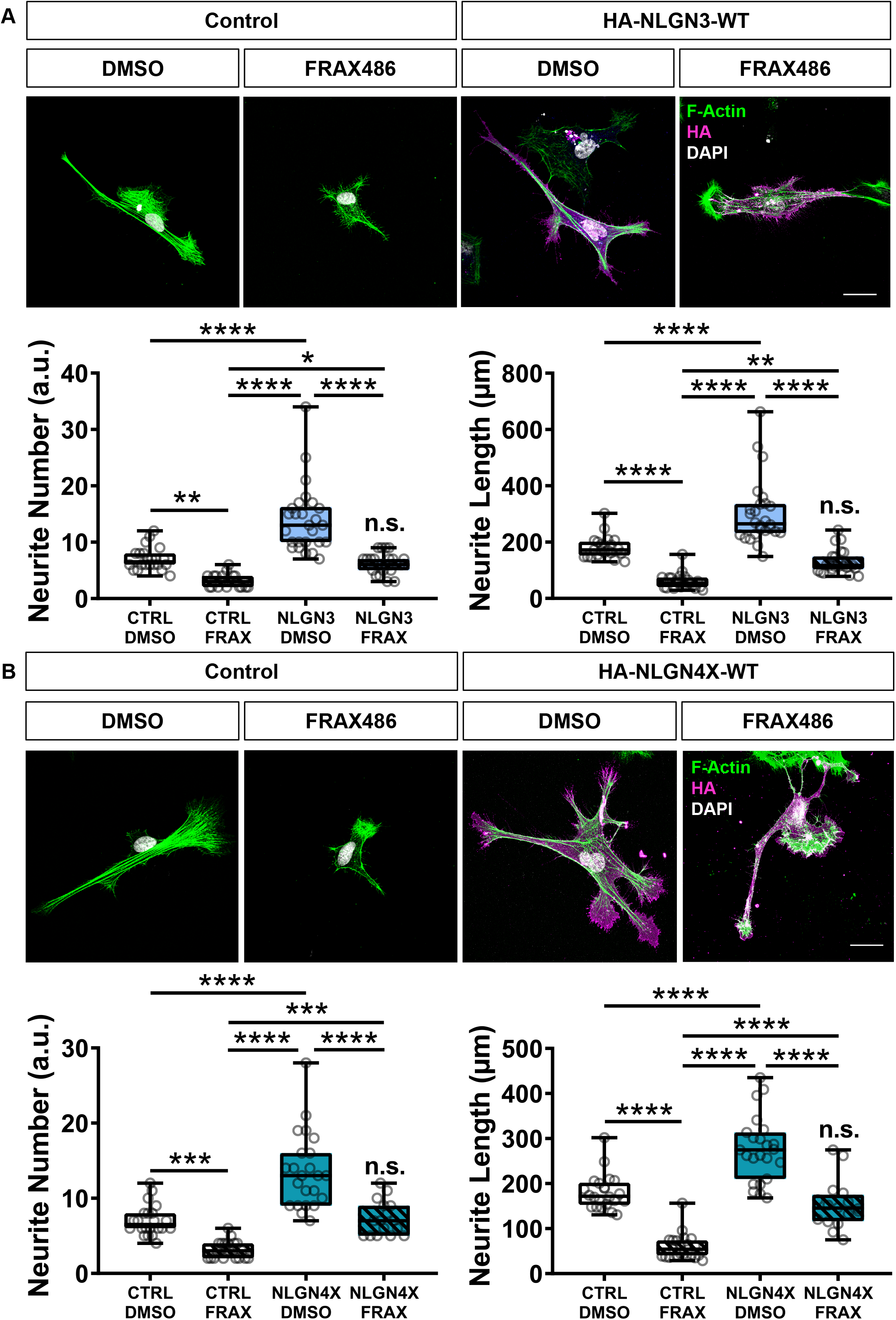
Increases in neuritogenesis induced by ectopic NLGN3/4X-WT expression are attenuated by PAK1 inhibition. (A) Representative images and data showing ectopic NLGN3-WT mediated increases in neurite number and length are attenuated by PAK1 inhibition. Scale bar = 25µm. Number: Kruskal-Wallis: χ2(4)=69.19, p<0.0001; CTRL-DMSO/CTRL-FRAX - Dunn: Mean rank difference: 34.30, p=0.0001, n=25; CTRL-DMSO/NLGN3-DMSO - Dunn: Mean rank difference: −30.03, p=0.0007, n=25; CTRL-DMSO/NLGN3-FRAX - Dunn: Mean rank difference: 5.83, p>0.99, n=25. Length: Kruskal-Wallis: χ2(4)=74.33, p<0.0001; CTRL-DMSO/CTRL-FRAX - Dunn: Mean rank difference: 41.77, p<0.0001, n=25; CTRL-DMSO/NLGN3-DMSO - Dunn: Mean rank difference: −32.40, p=0.017, n=25; CTRL-DMSO/NLGN3-FRAX - Dunn: Mean rank difference: 17.38, p=0.19, n=25. (B) Representative images and data showing ectopic NLGN4X-WT mediated increases in neurite number and length are attenuated by PAK1 inhibition. Scale bar = 25µm. Number: Kruskal-Wallis: χ2(4)=62.30, p<0.0001; CTRL-DMSO/CTRL-FRAX - Dunn: Mean rank difference: 29.75, p=0.0002, n=25; CTRL-DMSO/NLGN4X-DMSO - Dunn: Mean rank difference: −26.75, p=0.001, n=25; CTRL-DMSO/NLGN4X-FRAX - Dunn: Mean rank difference: −1.27, p>0.99, n=25. Length: Kruskal-Wallis: χ2(4)=62.40, p<0.0001; CTRL-DMSO/CTRL-FRAX - Dunn: Mean rank difference: 35.09, p<0.0001, n=25; CTRL-DMSO/NLGN4X-DMSO - Dunn: Mean rank difference: −20.85, p=0.02, n=25; CTRL-DMSO/NLGN4X-FRAX - Dunn: Mean rank difference: 10.25, p>0.99 n=25.

## Discussion

CAMs (cell-adhesion molecules) are key regulators of neuritogenesis, particularly at the growth cone leading edge^37^. Gene mutations in the trans-synaptic neurexin-neuroligin cell-adhesion complex are frequently associated with ASD pathogenesis, particularly neuroligin-3 and neuroligin-4X (NLGN3/4X)^13^. However, the role of NLGN3/4X during human cellular neurodevelopment is unknown. Furthermore, the functional consequences of ASC-associated NLGN3/4X mutations have yet to be investigated in human neurodevelopment. Here we demonstrate novel roles for NLGN3/4X in early human neurodevelopment. Specifically, we demonstrate NLGN3 and NLGN4X nanodomains promote neuritogenesis during cellular neurodevelopment via actin filament organization within the growth cone, mediated by PAK1 signaling. We also observed that growth cones of immature neurons ectopically expressing NLGN3/4X-WT cotreated with the PAK1 phosphorylation inhibitor, FRAX486, appeared similar in size and structure to growth cones ectopically expressing NLGN3-R451C or NLGN4X-D396. Furthermore, we demonstrate a novel feedback loop between PAK1 and NLGN3/4X nanodomain clustering which contributes to this growth cone enlargement and, by extension, neuritogenesis. Our data reveal novel roles for both NLGN3 and NLGN4X in the development of human cortical neurons, which is not replicated by ASD-associated mutants of these adhesion proteins **(Figure 8)**.

**Figure 8.**
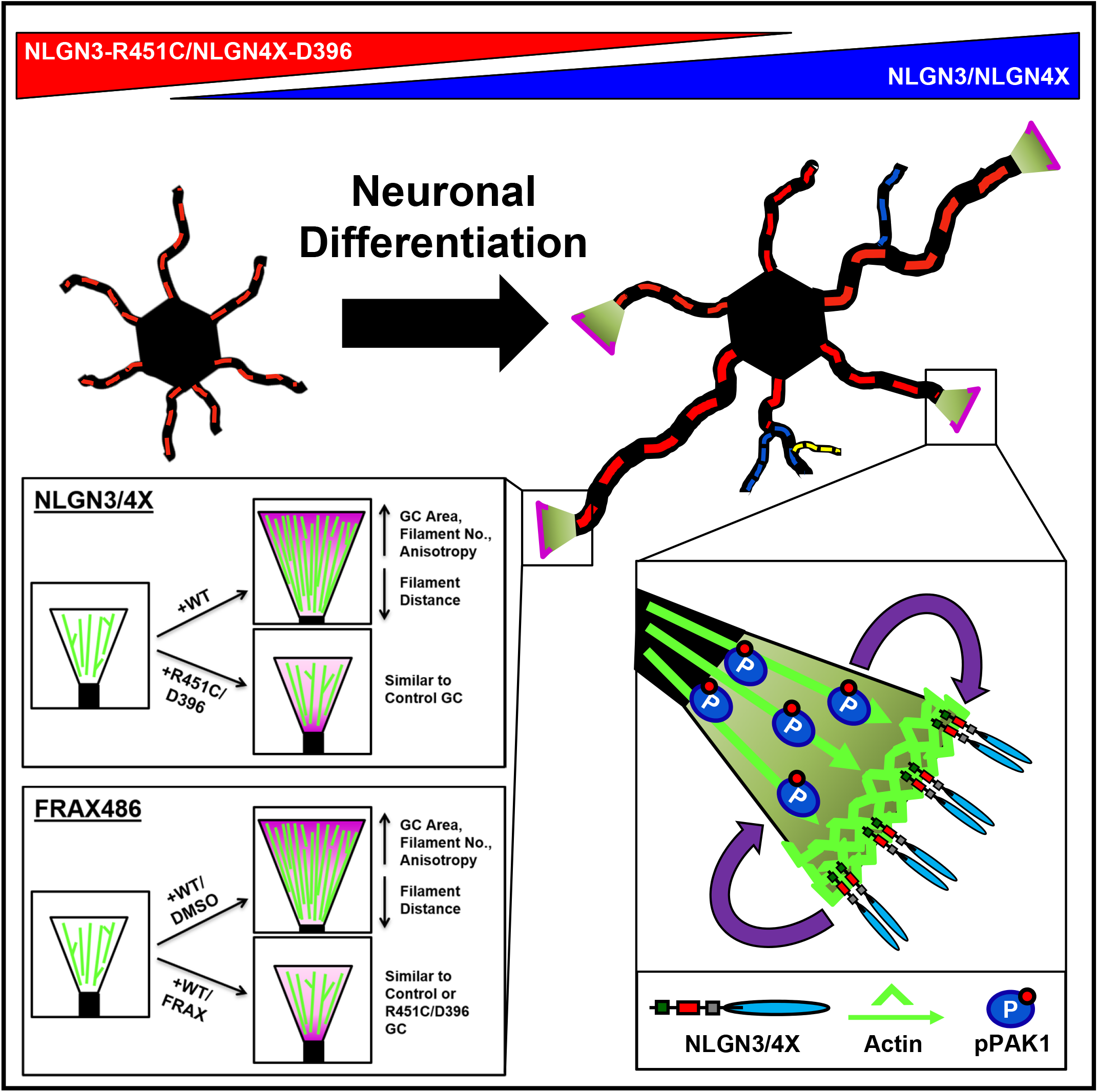
– Diagram summarizing all primary findings of this research including the effects on neuritogenesis (upper), the effects on growth cone actin (lower left), and the molecular mechanism involved in both phenotypes (lower right). NLGN, neuroligin; GC, growth cone; WT, wild type; DMSO, dimethyl sulfoxide; FRAX, FRAX486; PAK1, p21-activated kinase.

Broadly, our data indicates that WT NLGN3 and NLGN4X exert similar effects on the development of neuronal morphology. Similar increases in neurite length/number, growth cone area, and actin filament metrics were observed between the two proteins. Critically, both NLGN3 and NLGN4X demonstrated an ability to regulate PAK1 and cofilin activity, indicating that these CAMs signal via the same pathway to regulate the actin cytoskeleton. The similarity in effect and signaling pathways activated by these proteins is consistent with the comparatively high amino acid sequence homology between NLGN3 and NLGN4X (69.026% homology)^38^,^39^ and the finding that NLGN4 evolved rapidly from other NLGN gene sequences in mice^40^. Additionally, NLGN4X overexpression was recently found to rescue aberrant neurite outgrowth induced by ZNF804A knockdown^41^. Combined, these suggest necessary roles for both proteins in mammalian neuritogenesis as well as a degree of functional overlap between NLGN3 and NLGN4X. However, it is unlikely that these proteins operate alone. Accumulating evidence has linked NLGNs with other proteins involved in dendrite morphogenesis, such as Rap. Rap proteins are a subfamily of the Ras superfamily of small GTPases. Small GTPases have been extensively implicated in neuritogenesis and as signaling mediators for CAMs^1, 2^. NLGN3 has previously been shown to interact with the Rap guanine-nucleotide exchange factor, Epac2 (RapGEF4) thus controlling Rap signaling^42^. Rare variants in Epac2 have been associated with ASC and have been implicated in impaired basal dendrite morphogenesis^43^. It is also noteworthy that NLGN1 can regulate Rap activity via the activation of spine-associated Rap GTPase (SPAR) resulting in the modulation of LIMK/cofilin-mediated actin organization^44^. The importance for the correct regulation of such a pathway is highlighted by increasing evidence that ASC-associated mutations in SynGAP, a GAP for Rap, play a significant role in the development of neuronal and synaptic morphology^45^. Combined, these findings suggest several ASC-associated proteins operate via different signaling pathways prior to PAK1/cofilin but result in a similar modulation of actin dynamics at the endpoint. Furthermore, this suggests the ability of NLGN3/4X to regulate the actin-cytoskeleton via PAK1/cofilin is likely mediated by specific binding partners.

NLGNs binding partner NRXN has also been implicated in dendritogenesis. The NRXN-NLGN interaction is of particular importance as questions remain about exactly if/how NLGN3/4X are activated extracellularly to induce growth cone enlargement and neuritogenesis. For example, it is unclear if the growth cone enlarging effects of NLGN3/4X are activated by cis-or trans-binding mechanisms given the canonical pre/post-synaptic NRXN-NLGN configuration is not the focus of this research. It was recently discovered that NRXN-NLGN interactions play a key role in sexually dimorphic neurite plasticity during *Caenorhabditis elegans* cellular neurodevelopment^11^. Additionally, NRXN1β was found to induce neurite outgrowth in HEK293 cells and rat primary hippocampal neurons via an interaction with NLGN1^46^. Lastly, interactions between NLGN1 and NRXN were found to extensively modulate dendritogenesis by stabilizing filopodia in *Xenopus* and *Drosophila* neurodevelopment^9, 17^. These multiple lines of evidence indicate NRXN-NLGN interactions are likely a critical component of neurodevelopment which is evolutionarily conserved from Drosophila to humans. Combined with the evidence discussed previously, this suggests NLGNs may operate as components of a larger macromolecular structure with other proteins often associated with ASC, such as Epac2, SynGAP, or NRXN1 to regulate (or dysregulate) cytoskeletal remodeling during neurodevelopment and in the mature brain. However, many of these ASC-associated proteins may influence the cytoskeleton via the common mechanism of PAK1 but by different routes.

Despite the functionally compromising nature of the ASC-associated mutations, few dominant negative effects were observed when either mutant variant was ectopically expressed. This suggests the mutant proteins are not noticeably cytotoxic. Rather, the mutant proteins lack of appropriate localization and clustering at the growth cone leading edge resulted in an inability to gain function rather than mimic a loss of function i.e. growth cones and neurites do not shrink as a result of ectopic NLGN3-R451C/NLGN4X-D396 expression but also do not expand or extend completely as intended. It is noteworthy that we have utilized an ectopic-based “gain of function” approach to investigate the role of NLGN3/4 and the impact of ASC-mutations on their function. One potential caveat of this approach could arise due to a compensatory mechanism induced by endogenous expression of NLGN3/4X. Indeed, previous studies have shown that up or down regulation of NLGNs can produce conflicting results, depending on the cellular context^32, 47, 48^. Alterations in the expression of NLGNs and their subsequent impact on their cellular function may, therefore, be obfuscated by other NLGNs. This complication may further obfuscate subtle effects of ASC-associated mutations due to endogenous NLGN3/4X expression or upregulation of other NLGNs to compensate for the loss of function. This notion is particularly highlighted by the lack of effect of NLGN triple-KO on synapses^14^, as well as potential off target effects on the cytoskeleton induced by shRNA approaches^49^. Hence, we have opted to use a gain of function approach to disentangle NLGN3/4X function due to the emerging complex nature of endogenous NLGN interference.

While no dominant negative effects were found, subtle differences between NLGN3/4X-WT and their mutant variants were observed. For example, a significant difference in anisotropy was found between control and NLGN3-WT growth cone filaments. However, no significant differences in anisotropy were observed for NLGN4X-WT. This suggests NLGN4X has a subtly different function to NLGN3 in growth cones, perhaps suggesting NLGN3 has more of a stabilizing effect on actin filaments while NLGN4X has a destabilizing or less stabilizing effect. Additionally, a significant dominant negative effect was observed for filament distance between control and NLGN4X-D396. This was not observed for the NLGN3-R451C mutation. This suggests the NLGN4X-D396 mutation does have a deleterious effect on growth cones, but the observable effects may be diluted by the aforementioned compensatory mechanisms. These differences in actin filament organization would not have been observable without super-resolution microscopy and highlights the importance of using this advanced microscopy technique to further our understanding of subcellular molecular interactions. Lastly, the significance patterns in PAK and cofilin activation were slightly different between control, NLGN3/4X-WT, and their mutant variants. No dominant negative effects were observed, however, a significant increase in cofilin activation was observed between control and NLGN3-R451C, although not as substantial of an increase as NLGN3-WT. This partial increase in cofilin activity is likely due to the remaining partial functionality of the NLGN3-R451C protein. This was not observed in the NLGN4X-D396 condition, likely due to the complete loss of function of the NLGN4X-D396 protein.

The novel finding of a NLGN-PAK1 feedback loop dependent on molecular clustering at the cell membrane was an intriguing discovery and not without precedence. Indeed, filamin, an actin cross-linking protein, was found to operate in a similar bidirectional loop with PAK1 to locally influence actin cytoskeletal dynamics and membrane ruffles^50, 51^. Furthermore, nanoscale clustering of the chemokine receptor CXCR4 at the leading edge of Jurkat T lymphocyte cells was found to promote dynamic actin rearrangement resulting in increased cell migration^52^. Evidence from mature neurons also suggests orchestrated nanoscale recruitment and clustering of cell-adhesion molecules (CAM) at synapses is critical to synaptic function^32^. For example, NLGN1 and N-cadherin were found to cooperate to form clusters at synapses in mature hippocampal neurons, ultimately promoting synapse formation^53^. This synaptic clustering function was also demonstrated for NLGN3. Mature hippocampal neurons treated with purified Wnt3a exhibited increased NLGN3 recruitment to dendritic processes and, ultimately, increased clustering of NLGN3 with PSD-95 at synapses^30^. Lastly, but perhaps most relevantly, transient clustering interactions between flowing actin filaments and immobilized N-cadherin/catenin complexes were demonstrated in neuronal growth cones^8^. This led to a local reduction of actin retrograde flow and increased growth cone migration. Combined, this evidence suggests the actin cytoskeleton responds dynamically to nanoscale clustering of specific molecules at the cell membrane. This response is likely to be similar in growth cones given the high dynamicity of growth cone actin and the integral role of cell adhesion molecules in the growth cone, therefore, this system may also be a component of neuritogenesis during cellular neurodevelopment. Perturbation of this nanoscale clustering via ASC-associated NLGN mutations may therefore lead to alterations in growth cone motility, ultimately contributing to subtly atypical neurodevelopment. Additionally, the similarity in structure of NLGN3/4X-WT and FRAX486 cotreated growth cones to NLGN3/4X mutant growth cones suggests a link between the molecular mechanism discovered herein and the underlying mechanism that may be involved in the impaired cellular phenotypes seen when the ASC-associated mutant variants are ectopically expressed.

In summary, we demonstrate a novel function for NLGN3/4X during early human neurodevelopment in neuronal growth cones, particularly at the growth cone leading edge. The functional impact of this role at the growth cone was to significantly promote neuritogenesis in immature human neurons. We also illustrate the consequences of this new role on the actin cytoskeleton in that NLGN3/4X clustering has profound effects on growth cone F-actin remodeling, ultimately leading to significant changes in growth cone size and structure. These functional roles were found to be impaired by ASC-associated mutant forms of NLGN3/4X for which the clustering function was severely impaired. Furthermore, we show a link between NLGN3/4X clustering and PAK1 activation in the growth cone. Ultimately, leading to the discovery of a mechanistic feedback loop between NLGN3/4X and PAK1 in growth cones which drives actin dynamics, growth cone enlargement, and neuritogenesis.

## Methods

### Antibodies and Plasmids

Antibodies: mouse anti-HA monoclonal (BioLegend, 901503, 1:1000), rabbit anti-RFP polyclonal (MBL, PM005, 1:500), chicken anti-GFP polyclonal (Abcam, ab13970, 1:1000), chicken anti-Tuj1 polyclonal (Abcam, ab41489, 1:500), ActinGreen 488 ReadyProbe (Thermo Fisher, R37110, per manufacturer’s instructions), and rabbit anti-phospho-PAK1 (Ser144)/PAK2 (Ser141) (Cell Signaling Technology, 2606, 1:100). Alexa Fluor 488, 568, 633 (Life Technologies, 1:500) and 4’,6-diamidino-2-phenylindole (DAPI – D1306, Life Technologies, 1:50000) fluorescent secondary antibodies were used in all immunocytochemistry experiments where applicable. Immunoblotting: mouse anti-NLGN3 monoclonal (StressMarq, SMC-471D, 1:1000), rabbit anti-NLGN3 polyclonal (Synaptic Systems, 129113, 1:1000), rabbit anti-HA polyclonal (ProteinTech, 51064-2-AP, 1:1000), rabbit anti-NLGN4X monoclonal (Abcam, ab181251, 1:1000), mouse anti-total cofilin monoclonal (ProteinTech, 66057-1-lg, 1:3000), rabbit anti-phospho-cofilin (Ser3) polyclonal (Cell Signaling Technology, 3311, 1:1000), rabbit anti-total PAK1/2/3 (Cell Signaling Technology, 2604), rabbit anti-phospho-PAK1 (Ser144)/PAK2 (Ser141) (Cell Signaling Technology, 2606, 1:1000), rabbit anti-HRP (Life Technologies, G-21234, 1:10000), mouse anti-HRP (Life Technologies, A16078, 1:10000) **(Supplemental Table 1)**.

Cloned HA-tagged NLGN3-WT, NLGN3-R451C, NLGN4X-WT and NLGN4X-D396 plasmids were gifts from Prof. Peter Scheiffele^23^. A pmCherry-N1 plasmid (ClonTech, 632523) was utilised as a morphological marker in HEK293 immunocytochemistry experiments. A peGFP-N2 plasmid (ClonTech, 632483) was utilised as a morphological marker in subsequent proliferating CTX0E16 experiments due to red fluorescent protein aggregation in CTX0E16 hNPCs.

### Cell Culture, Transfection, and Replication

Antibody validation, protrusion outgrowth, and PAK/cofilin mechanism experiments used human embryonic kidney cells (HEK293) cultured in DMEM:F12 (Sigma, D6421) supplemented with 10% foetal bovine serum (ClonTech, 631107) and 1% L-glutamine (Sigma, G7513), in a 37°C/5% CO2 atmosphere. Cells were seeded at 30-40% confluence on acid-washed glass coverslips 24 hours before transfection. For antibody validation, three biological replicate whole cell lysate samples were generated (see immunoblotting section) and separated per lane. One biological replicate was considered as a single passage/plating between flasks of fully confluent cells. For PAK/cofilin mechanism experiments, three biological replicate whole cell lysate samples per condition (control, wildtype NLGN, and mutant NLGN) were generated 48 hours post-transfection. The protrusion outgrowth experiment was conducted in HEK293 cells once to confirm the outgrowth phenotype.

The conditionally immortalised cortically derived human neural progenitor cell line (hNPC) CTX0E16 was obtained from ReNeuron Ltd. (Guildford, UK) under a Material Transfer Agreement. CTX0E16 hNPCs were derived from 12-week foetal cortical neuroepithelium and conditionally immortalized using a c-mycER^TAM^ transgene. Characterization of the CTX0E16 cell line is described in detail elsewhere^54, 55^. Proliferation and neuralization of CTX0E16 cells were carried out as previously described^54^. Briefly, CTX0E16 cells were neuralized by replacing 4-hydroxytamoxifen (Sigma, H7904) supplemented DMEM:F12 medium (Sigma, D6421) with Neurobasal medium (Invitrogen, 12348017) supplemented with serum-free B27 (Life Technologies, 17504044).

For endogenous NLGN3/4X expression experiments, three biological replicate whole cell protein or RNA lysates per time point (NPC and neuron) were generated in parallel (see immunoblotting or RNA isolation sections). Super-resolution images of endogenous NLGN3/4X expression were conducted once to assess endogenous protein localisation in immature neurons.

All transfections in HEK293 and CTX0E16 hNPCs were carried out using Lipofectamine 2000 (Invitrogen, 17504044), per the manufacturer’s instructions. Briefly, 2 µg of each HA-NLGN and 2 µg of peGFP-N2 or mCherry (where applicable) construct were mixed with 2 µl of Lipofectamine 2000 in 100 µl DMEM:F12 and incubated for 20 minutes in a 37°C/5% CO2 atmosphere. The DNA:Lipofectamine 2000 mixture was added dropwise to HEK293 or CTX0E16 hNPCs which were then incubated for 4 hours at 37°C, before being transferred to new wells containing fresh media. Proliferating HEK293 or CTX0E16 hNPCs recovered for 48 hours in an incubator at 37°C/5% CO2 prior to fixation and immunocytochemistry. Differentiating CTX0E16 hNPCs recovered for 48 hours and continued differentiating for a further 24 hours in an incubator at 37°C/5% CO2 prior to fixation and immunocytochemistry.

Super-resolution assessment of nanodomains was conducted once to confirm their presence in and at the leading edge of growth cones. For neurite outgrowth experiments, five images per condition (control, wildtype NLGN, mutant NLGN) were acquired and quantified for each of three biological replicates. One biological replicate was considered as a single passage/plating between flasks of fully confluent cells. Line scan analyses of exogenously expressed growth cone and neurite NLGN3/4X were conducted once to demonstrate the localisation disparity between wildtype and mutant NLGN. For super-resolution growth cone actin analyses, five images per condition (control, wildtype NLGN, mutant NLGN) were acquired and quantified for each of three biological replicates (see microscopy and quantification sections). For super-resolution analysis of growth cone pPAK1 and wildtype NLGNs, five images per condition (control, wildtype NLGN) were acquired and quantified for each of three biological replicates.

The brain penetrant and orally bioavailable p21-activated kinase (PAK) inhibitor FRAX486 (Tocris, 5190) was used to inhibit PAK1-4 in pharmacological inhibition/rescue experiments. FRAX486 was diluted in dimethyl sulphate (DMSO) to 10 mM per manufacturer’s instructions. Differentiating hNPCs were treated with FRAX486 diluted in differentiation media 24 hours after transfection with HA-NLGN3/4X constructs at a final concentration of 50 nM^35, 36^. DMSO was used as a vehicle treatment for control at the same dilution. FRAX486 and vehicle treated hNPCs continued differentiating for a further 48 hours in an incubator at 37°C/5% CO2 prior to fixation and immunocytochemistry. For super-resolution growth cone FRAX486 attenuation experiments, five images per condition (control DMSO, control FRAX, NLGN DMSO, NLGN FRAX) were acquired and quantified for each of three biological replicates. For FRAX486 treated growth cone NLGN cluster experiments, five images per condition (control DMSO, control FRAX, NLGN DMSO, NLGN FRAX) were acquired and quantified for each of three biological replicates. For neurite FRAX486 attenuation experiments, five images per condition (control DMSO, control FRAX, NLGN DMSO, NLGN FRAX) were acquired and quantified for each of three biological replicates.

### Microscopy

Representative images of preliminary experiments in HEK293 and CTX0E16 cells, proliferating CTX0E16 experiments and all images for data analysis were acquired using a Zeiss Axio Imager Z1 epifluorescent microscope with ApoTome attachment using a 40x oil-immersion objective (Carl Zeiss AG). Representative images of further experiments investigating neurite outgrowth in differentiated CTX0E16 cells were acquired using a Leica SP-5 confocal microscope with a 100x oil-immersion objective (Leica microsystems). Z-stacks of 5 individual cells per condition across three biological replicates were obtained for statistical data analysis. 10 images were taken per Z-stack at a slice distance of 0.5 μm.

Super-resolved images of growth cones were collected using a Visitech-iSIM module coupled to a Nikon Ti-E microscope with a Nikon 100x 1.49 NA TIRF oil immersion lens (Nikon, Japan). Blue fluorescence was excited with a 405 nm laser and emission filtered through a 460/50 filter. Green fluorescence was excited with a 488 nm laser and emission filtered through a 525/50 filter. Red Fluorescence was excited with a 561 nm laser and emission filtered through a 630/30 filter. Far Red fluorescence was excited with a 640 nm laser and emission filtered through a 710/60 filter. Multiple images at focal planes were collected spaced apart by 0.05 μm. Data were deconvolved using a Richardson-Lucy algorithm specific to the iSIM mode of imaging to increase contrast and resolution using the supplied NIS-Elements Advanced Research software (Nikon, Japan, v4.6).

### Immunoblotting

Cultured cells were lysed in radioimmunoprecipitation lysis buffer consisting of 20 mM Tris; pH 7.2, 150 mM NaCl, 1% Triton-X-100, 5 mM EDTA; pH 8, 0.1% SDS, 1% sodium deoxycholate with additional phosphatase inhibitors (Sigma, P0044). Detergent soluble lysates were sonicated and centrifuged to remove cell debris. Samples were resolved by SDS-PAGE, transferred to a nitrocellulose or PVDF membrane and blocked for 1 hour in 5% bovine serum albumin (Sigma, A7906) in TBS-T. Membranes were then immunoblotted with primary antibodies overnight at 4°C, followed by incubation with anti-mouse or anti-rabbit horseradish peroxidase (HRP) conjugated secondary antibodies for 1 hour at room temperature. Membranes were then incubated in Clarity electrochemiluminescence substrate (Bio-Rad, 1705061) for 5 minutes and subsequently scanned using the Bio-Rad ChemiDoc MP (Bio-Rad). Band intensity was quantified by densitometry using Image Lab software (Bio-Rad, v6.0).

### RNA Isolation, cDNA Synthesis, RT-PCR, and qPCR

Proliferating hNPCs (n=3) or differentiating immature neurons (n=3) CTX0E16s were pelleted and lysed in TRI Reagent (Ambion, AM9738). Total RNA was then extracted per the manufacturer’s instructions. Residual genomic DNA was removed from each of 6 biological replicates using the TURBO DNA-*free*™ Kit (Life Technologies, AM1907) per the manufacturer’s instructions. cDNA was synthesized from 1 μg of total RNA from each extraction using random decamers (Ambion, AM5722G) and SuperScript III Reverse Transcriptase (Invitrogen, 18080044), per the manufacturer’s instructions.

To determine the expression of specific genes, primers were designed to target all known RefSeq transcripts of genes of interest, sourced from the UCSC Genome Browser website (http://genome.ucsc.edu) **(Supplemental Table 2)**. Primers were designed to span intronic regions of the selected genes to ensure specific amplification of mRNA, even in the presence of DNA contamination. Reactions were carried out in a total volume of 20 µl containing diluted cDNA, 1 X HOT FIREPol Blend Master Mix (Solis Biodyne, 04-25-00125) and primers at 200 nM, using a GS4 thermal cycler. Samples were separated and visualised by agarose gel electrophoresis.

For quantitative expression analysis, 20 µl cDNA samples from SuperScript III reactions were diluted with a further 120 µl of nuclease-free H2O. Reactions were carried out in a total volume of 20 µl, containing diluted cDNA, 1× HOT FIREPol® EvaGreen® q-PCR Mix (Solis Biodyne, 08-25-00001) and primers at 200 nM, using an MJ Research Chromo 4 (Bio-Rad) and MJ Opticon Monitor analytic software (Bio-Rad). Triplicate qPCR reactions were performed to measure each gene in each cDNA sample. The level of each gene was measured against a standard curve constructed by serial dilution of pooled cDNA from all assayed samples. A relative value was thus obtained for each of the three triplicate reactions for each cDNA sample. Mean measures of target genes were then normalized against a geometric mean determined from 2 internal control genes (ALG2 & RPL6) for each cDNA sample to yield a relative target gene expression value for all samples. ALG2 & RPL6 were identified as suitable internal controls based on a combination of previous whole-genome microarray data of CTX0E16 cells, where it showed the least variability (in terms of standard deviation) across conditions and a housekeeper screen qPCR. Normalized qPCR target gene expression values were compared between hNPC or immature neuron CTX0E16 cells.

### Quantification and Statistical Analysis

All neurite outgrowth images were quantified using the NeuronJ ImageJ plug-in (http://www.imagescience.org/meijering/software/neuronj/ v1.4.2) which allowed for manual tracing and labelling (primary, secondary, or tertiary) of individual neurites^56^ **(Supplemental Figure 7A)**. Neurite inclusion/exclusion criteria were established visually while tracing and later re-established once all neurites were traced via a <3 μm neurite length exclusion threshold set via logic test in MS Excel. 5 cells per condition were quantified across three biological replicates.

Growth cone area was quantified using NIS-Elements Advanced Research software (Nikon, Japan). Growth cone filament number was quantified using line scan analysis in ImageJ **(Supplemental Figure 7B & C)**. Growth cone filament skeletons were generated using the ImageJ plug-in Ridge Detection^57^. Filament length was also quantified using Ridge Detection. Growth cone filament anisotropy was quantified using the ImageJ plug-in FibrilTool^58^.

All immunoblotting and RT-qPCR data were processed post-hoc in MS Excel to eliminate batch effects^59^. Briefly, the sum of all conditions in each data set was calculated. Each condition was then divided by the sum of all conditions to remove batch effects. Data were then log transformed and plotted. All datasets were tested for normality using the D’Agostino & Pearson normality test prior to inferential statistical analyses ^60^. Datasets found to be normally distributed were analysed using parametric statistical tests while datasets found to be abnormally distributed were analysed using their non-parametric equivalent. Two-tailed unpaired students t-test or Mann Whitney U test was used for endogenous expression protein/RNA analyses with an alpha level of 0.05. Ordinary one-way analysis of variance (ANOVA) with Bonferroni correction for multiple comparisons or Kruskal-Wallis test with Dunn’s post-hoc correction was used for total neurite outgrowth analyses with an alpha level of 0.05 (MRD, Mean Rank Difference). Two-way ANOVA with Bonferroni correction for multiple comparisons was used for primary, secondary, and tertiary neurite outgrowth analyses. Ordinary one-way ANOVA with Bonferroni correction for multiple comparisons or Kruskal-Wallis test with Dunn’s post-hoc correction was used for all super-resolution growth cone actin analyses, including FRAX486 attenuation analyses with an alpha level of 0.05. Two-tailed unpaired students t-test or Mann Whitney U test was used for FRAX486 treated growth cone NLGN cluster analyses. Lastly, ordinary one-way ANOVA with Bonferroni correction for multiple comparisons or Kruskal-Wallis test with Dunn’s post-hoc correction was used for FRAX486 treated neurite outgrowth analyses. All data visualisations were generated in GraphPad Prism 7.0 (GraphPad Software, La Jolla California USA, http://www.graphpad.com/scientific-software/prism/). All data are shown as mean ± standard error of the mean (SEM) to two decimal places where necessary and all error bars represent SEM.

## Data Availability

All data supporting the findings of this manuscript are available from the corresponding authors upon reasonable request. NLGN3/4X mRNA expression data are freely available from the BrainSpan Atlas of the Developing Human Brain compiled primarily by the Allen Institute for Brain Science.

## Contributions

Conceptualization, D.P.S.; Methodology, D.P.S. and N.J.F.G.; Investigation, N.J.F.G., P.J.M.D., R.R.R.D., G.C., and P.R.; Writing – Original Draft, N.J.F.G.; Writing – Review & Editing, D.P.S. and N.J.F.G.; Visualization, N.J.F.G.; Supervision, D.P.S.

## Acknowledgements

The authors gratefully thank Professor Philip Gordon-Weeks for his insightful editorial comments and feedback on the manuscript and figures. The study was supported by grants from the Wellcome Trust ISSF Grant (No. 097819) and the King’s Health Partners Research and Development Challenge Fund, a fund administered on behalf of King’s Health Partners by Guy’s and St Thomas’ Charity awarded to DPS; the Brain and Behavior Foundation (formally National Alliance for Research on Schizophrenia and Depression (NARSAD); Grant No. 25957), awarded to DPS; the European Autism Interventions (EU-AIMS), and the Innovative Medicines Initiative Joint Undertaking under grant agreement no. 115300, resources of which are composed of financial contribution from the European Union’s Seventh Framework Programme (FP7/2007-2013) and EFPIA companies’ in kind contribution (DPS). We thank the Wohl Cellular Imaging Centre (WCIC) at the IoPPN, Kings College, London, for help with microscopy.

## References

1. Jan Y-N, Jan LY. Branching out: mechanisms of dendritic arborization. Nature Reviews Neuroscience 11, 316 (2010).

2. Arimura N, Kaibuchi K. Neuronal polarity: from extracellular signals to intracellular mechanisms. Nature Reviews Neuroscience 8, 194 (2007).

3. Polleux F, Snider W. Initiating and growing an axon. Cold Spring Harbor perspectives in biology 2, a001925 (2010).

4. Lowery LA, Van Vactor D. The trip of the tip: understanding the growth cone machinery. Nature reviews Molecular cell biology 10, 332 (2009).

5. Pak CW, Flynn KC, Bamburg JR. Actin-binding proteins take the reins in growth cones. Nature Reviews Neuroscience 9, 136 (2008).

6. Delorme V, et al. Cofilin activity downstream of Pak1 regulates cell protrusion efficiency by organizing lamellipodium and lamella actin networks. Developmental cell 13, 646–662 (2007).

7. Flynn KC, et al. ADF/cofilin-mediated actin retrograde flow directs neurite formation in the developing brain. Neuron 76, 1091–1107 (2012).

8. Garcia M, Leduc C, Lagardère M, Argento A, Sibarita J-B, Thoumine O. Two-tiered coupling between flowing actin and immobilized N-cadherin/catenin complexes in neuronal growth cones. Proceedings of the National Academy of Sciences, 201423455 (2015).

9. Constance WD, et al. Neurexin and Neuroligin-based adhesion complexes drive axonal arborisation growth independent of synaptic activity. eLife 7, (2018).

10. Harkin LF, et al. Neurexins 1–3 each have a distinct pattern of expression in the early developing human cerebral cortex. Cerebral Cortex 27, 216–232 (2017).

11. Hart MP, Hobert O. Neurexin controls plasticity of a mature, sexually dimorphic neuron. Nature 553, 165 (2018).

12. Chubykin AA, et al. Activity-dependent validation of excitatory versus inhibitory synapses by neuroligin-1 versus neuroligin-2. Neuron 54, 919–931 (2007).

13. Südhof TC. Neuroligins and neurexins link synaptic function to cognitive disease. Nature 455, 903–911 (2008).

14. Varoqueaux F, et al. Neuroligins determine synapse maturation and function. Neuron 51, 741–754 (2006).

15. Clarris HJ, McKeown S, Key B. Expression of neurexin ligands, the neuroligins and the neurexophilins, in the developing and adult rodent olfactory bulb. International Journal of Developmental Biology 46, 649–652 (2004).

16. Paraoanu LE, Becker-Roeck M, Christ E, Layer PG. Expression patterns of neurexin-1 and neuroligins in brain and retina of the chick embryo: Neuroligin-3 is absent in retina. Neuroscience letters 395, 114–117 (2006).

17. Chen SX, Tari PK, She K, Haas K. Neurexin-neuroligin cell adhesion complexes contribute to synaptotropic dendritogenesis via growth stabilization mechanisms in vivo. Neuron 67, 967–983 (2010).

18. Gaugler T, et al. Most genetic risk for autism resides with common variation. Nature genetics 46, 881 (2014).

19. Basu SN, Kollu R, Banerjee-Basu S. AutDB: a gene reference resource for autism research. Nucleic Acids Research 37, D832–D836 (2009).

20. Kathuria A, et al. Stem cell-derived neurons from autistic individuals with SHANK3 mutation show morphogenetic abnormalities during early development. Mol Psychiatry 23, 735–746 (2018).

21. Schafer ST, et al. Pathological priming causes developmental gene network heterochronicity in autistic subject-derived neurons. 1 (2019).

22. Jamain S, et al. Mutations of the X-linked genes encoding neuroligins NLGN3 and NLGN4 are associated with autism. Nature genetics 34, 27–29 (2003).

23. Chih B, Afridi SK, Clark L, Scheiffele P. Disorder-associated mutations lead to functional inactivation of neuroligins. Human molecular genetics 13, 1471–1477 (2004).

24. Chanda S, Aoto J, Lee S-J, Wernig M, Südhof TCJMp. Pathogenic mechanism of an autism-associated neuroligin mutation involves altered AMPA-receptor trafficking. 21, 169 (2016).

25. Miller JA, et al. Transcriptional landscape of the prenatal human brain. Nature 508, 199–206 (2014).

26. Letellier M, et al. A unique intracellular tyrosine in neuroligin-1 regulates AMPA receptor recruitment during synapse differentiation and potentiation. Nature Communications 9, 3979 (2018).

27. Graf ER, Zhang X, Jin S-X, Linhoff MW, Craig AM. Neurexins induce differentiation of GABA and glutamate postsynaptic specializations via neuroligins. Cell 119, 1013–1026 (2004).

28. Boucard AA, Chubykin AA, Comoletti D, Taylor P, Südhof TC. A Splice Code for trans-Synaptic Cell Adhesion Mediated by Binding of Neuroligin 1 to α- and β-Neurexins. Neuron 48, 229–236 (2005).

29. Levinson JN, Li R, Kang R, Moukhles H, El-Husseini A, Bamji SX. Postsynaptic scaffolding molecules modulate the localization of neuroligins. Neuroscience 165, 782–793 (2010).

30. Medina MA, et al. Wnt/β-catenin signaling stimulates the expression and synaptic clustering of the autism-associated Neuroligin 3 gene. Translational psychiatry 8, 45 (2018).

31. Hoon M, et al. Neuroligin-4 is localized to glycinergic postsynapses and regulates inhibition in the retina. Proceedings of the National Academy of Sciences 108, 3053–3058 (2011).

32. Chamma I, Thoumine O. Dynamics, nanoscale organization, and function of synaptic adhesion molecules. Molecular and Cellular Neuroscience, (2018).

33. Haas KT, et al. Pre-post synaptic alignment through neuroligin-1 tunes synaptic transmission efficiency. eLife 7, e31755 (2018).

34. Edwards DC, Sanders LC, Bokoch GM, Gill GN. Activation of LIM-kinase by Pak1 couples Rac/Cdc42 GTPase signalling to actin cytoskeletal dynamics. Nature cell biology 1, 253 (1999).

35. Dolan BM, et al. Rescue of fragile X syndrome phenotypes in Fmr1 KO mice by the small-molecule PAK inhibitor FRAX486. Proceedings of the National Academy of Sciences of the United States of America 110, 5671–5676 (2013).

36. Hayashi-Takagi A, et al. PAKs inhibitors ameliorate schizophrenia-associated dendritic spine deterioration in vitro and in vivo during late adolescence. Proceedings of the National Academy of Sciences 111, 6461–6466 (2014).

37. Vitriol EA, Zheng JQ. Growth cone travel in space and time: the cellular ensemble of cytoskeleton, adhesion, and membrane. Neuron 73, 1068–1081 (2012).

38. Breuza L, et al. The UniProtKB guide to the human proteome. Database 2016, (2016).

39. Consortium U. UniProt: the universal protein knowledgebase. Nucleic acids research 45, D158–D169 (2016).

40. Bolliger MF, Pei J, Maxeiner S, Boucard AA, Grishin NV, Südhof TC. Unusually rapid evolution of Neuroligin-4 in mice. Proceedings of the National Academy of Sciences 105, 6421–6426 (2008).

41. Deans PM, et al. Psychosis risk candidate ZNF804A localizes to synapses and regulates neurite formation and dendritic spine structure. 82, 49–61 (2017).

42. Woolfrey KM, et al. Epac2 induces synapse remodeling and depression and its disease-associated forms alter spines. Nature neuroscience 12, 1275–1284 (2009).

43. Srivastava DP, et al. An autism-associated variant of Epac2 reveals a role for Ras/Epac2 signaling in controlling basal dendrite maintenance in mice. PLoS Biol 10, e1001350 (2012).

44. Liu A, et al. Neuroligin 1 regulates spines and synaptic plasticity via LIMK1/cofilin-mediated actin reorganization. The Journal of Cell Biology 212, 449 (2016).

45. Clement JP, et al. Pathogenic SYNGAP1 mutations impair cognitive development by disrupting maturation of dendritic spine synapses. Cell 151, 709–723 (2012).

46. Gjørlund MD, et al. Neuroligin-1 induces neurite outgrowth through interaction with neurexin-1β and activation of fibroblast growth factor receptor-1. The FASEB Journal 26, 4174–4186 (2012).

47. Chanda S, Hale WD, Zhang B, Wernig M, Südhof TC. Unique vs. Redundant Functions of Neuroligin Genes in Shaping Excitatory and Inhibitory Synapse Properties. Journal of Neuroscience, 0125–0117 (2017).

48. Biederer T, Kaeser PS, Blanpied TA. Transcellular nanoalignment of synaptic function. Neuron 96, 680–696 (2017).

49. Alvarez VA, Ridenour DA, Sabatini BL. Retraction of synapses and dendritic spines induced by off-target effects of RNA interference. Journal of Neuroscience 26, 7820–7825 (2006).

50. Vadlamudi RK, et al. Filamin is essential in actin cytoskeletal assembly mediated by p21-activated kinase 1. Nature cell biology 4, 681 (2002).

51. Rane CK, Minden A. P21 activated kinases: structure, regulation, and functions. Small GTPases 5, e28003 (2014).

52. Martínez-Muñoz L, et al. Separating Actin-Dependent Chemokine Receptor Nanoclustering from Dimerization Indicates a Role for Clustering in CXCR4 Signaling and Function. Molecular cell 70, 106–119. e110 (2018).

53. Aiga M, Levinson JN, Bamji SX. N-cadherin and neuroligins cooperate to regulate synapse formation in hippocampal cultures. Journal of Biological Chemistry 286, 851–858 (2011).

54. Anderson GW, et al. Characterisation of neurons derived from a cortical human neural stem cell line CTX0E16. Stem Cell Research Therapy 6, 149 (2015).

55. Pollock K, et al. A conditionally immortal clonal stem cell line from human cortical neuroepithelium for the treatment of ischemic stroke. Experimental neurology 199, 143–155 (2006).

56. Meijering E, Jacob M, Sarria JC, Steiner P, Hirling H, Unser M. Design and validation of a tool for neurite tracing and analysis in fluorescence microscopy images. Cytometry Part A 58, 167–176 (2004).

57. Steger C. An unbiased detector of curvilinear structures. IEEE Transactions on pattern analysis and machine intelligence 20, 113–125 (1998).

58. Boudaoud A, et al. FibrilTool, an ImageJ plug-in to quantify fibrillar structures in raw microscopy images. Nat Protocols 9, 457–463 (2014).

59. Degasperi A, Birtwistle MR, Volinsky N, Rauch J, Kolch W, Kholodenko BN. Evaluating strategies to normalise biological replicates of Western blot data. PloS one 9, e87293 (2014).

60. D’Agostino RB, Belanger A. A Suggestion for Using Powerful and Informative Tests of Normality. The American Statistician 44, 316–321 (1990).

