## Supplemental Information for "Nanoscopic Clustering of Neuroligin-3 and Neuroligin-4X Regulates Growth Cone Organization and Size"

**Supplemental Figures 1-7**

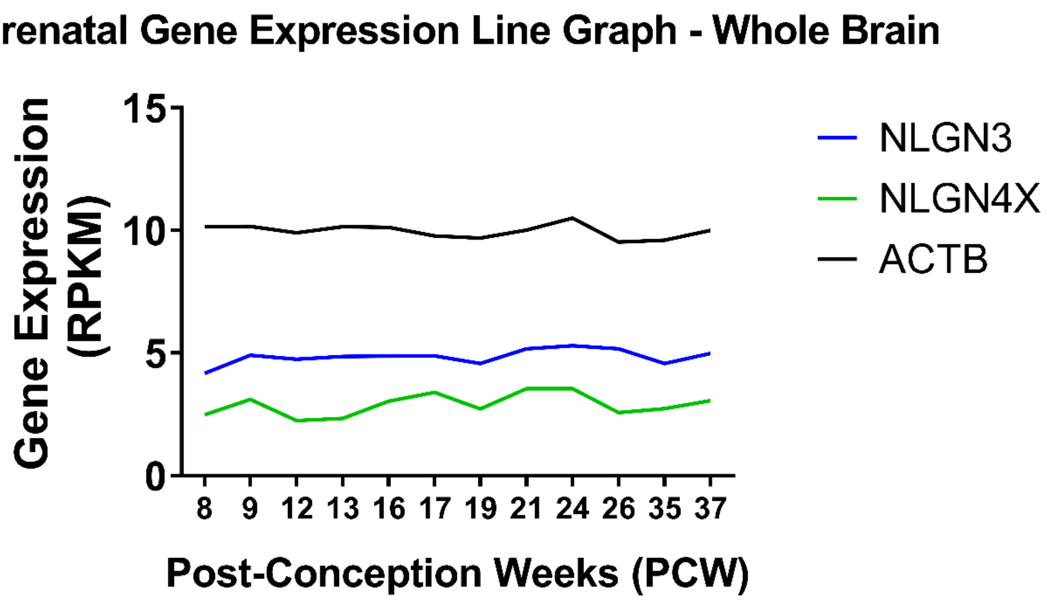


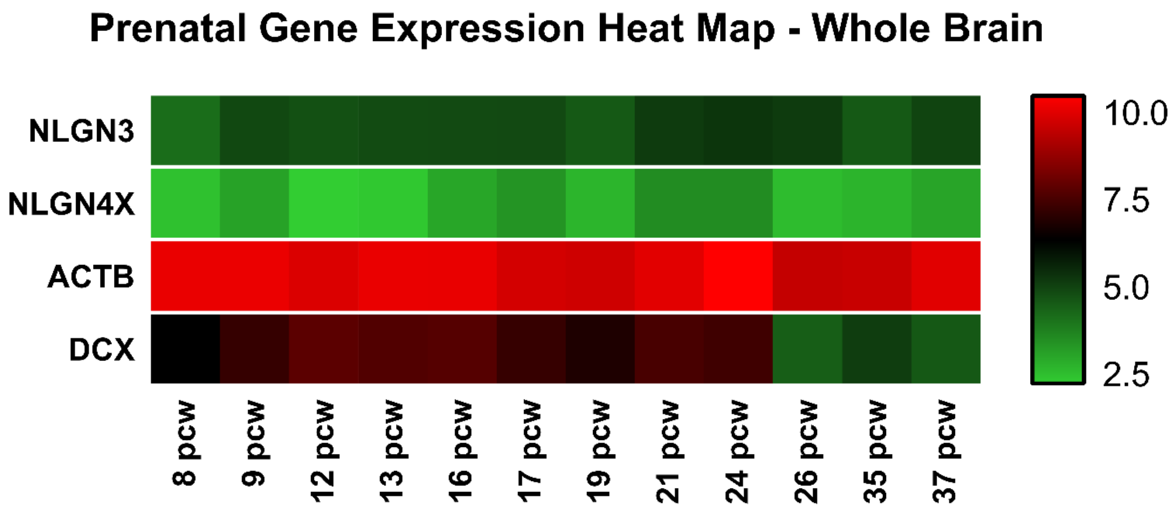


Gatford et al. Supplemental Figure 1

**Supplemental Figure 1** – Prenatal gene expression data reveals NLGN3 and NLGN4X are expressed during human prenatal neurodevelopment. (A) Prenatal human gene expression of NLGN3 and NLGN4X compared to beta-actin (ACTB) compiled from the Allen Brain Atlas shown as a line graph. (B) Prenatal human gene expression of NLGN3 and NLGN4X compared to beta-actin (ACTB) and doublecortin (DCX) compiled from the Allen Brain Atlas shown as a heat map.

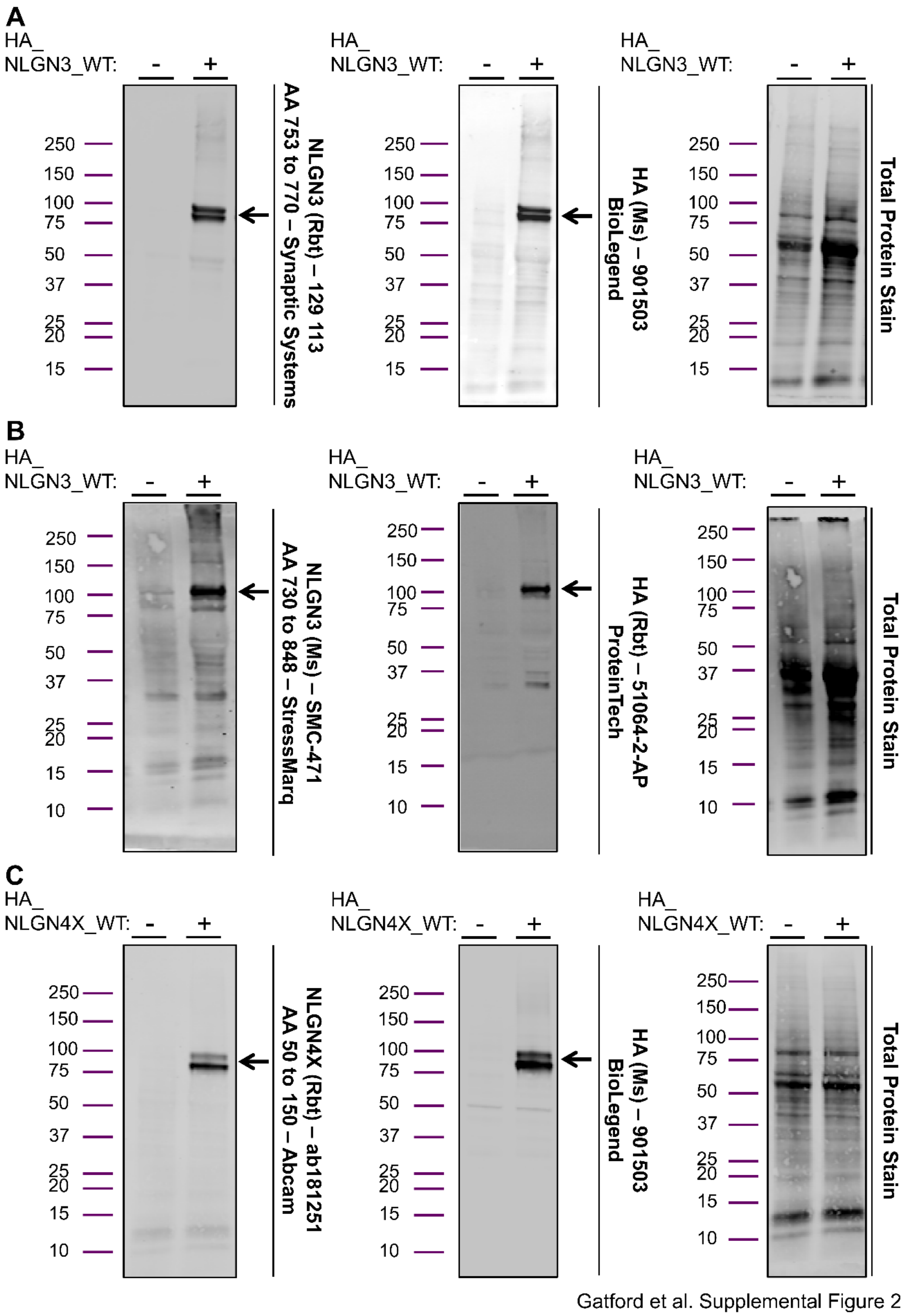


**Supplemental Figure 2** – NLGN3 and NLGN4X antibodies are highly specific (A) Representative western blots demonstrating the NLGN3 (Rabbit) antibody to NLGN3 AA 753 to 770 is highly specific to NLGN3 and the HA (Mouse) antibody is highly specific to HA. (B) Representative western blots demonstrating the NLGN3 (Mouse) antibody to NLGN3 AA 730 to 848 is also highly specific to NLGN3 and the HA (Rabbit) antibody is highly specific to HA. (C) Representative western blots demonstrating the NLGN4X (Rabbit) antibody to NLGN4X AA 50 150 is highly specific to NLGN4X.

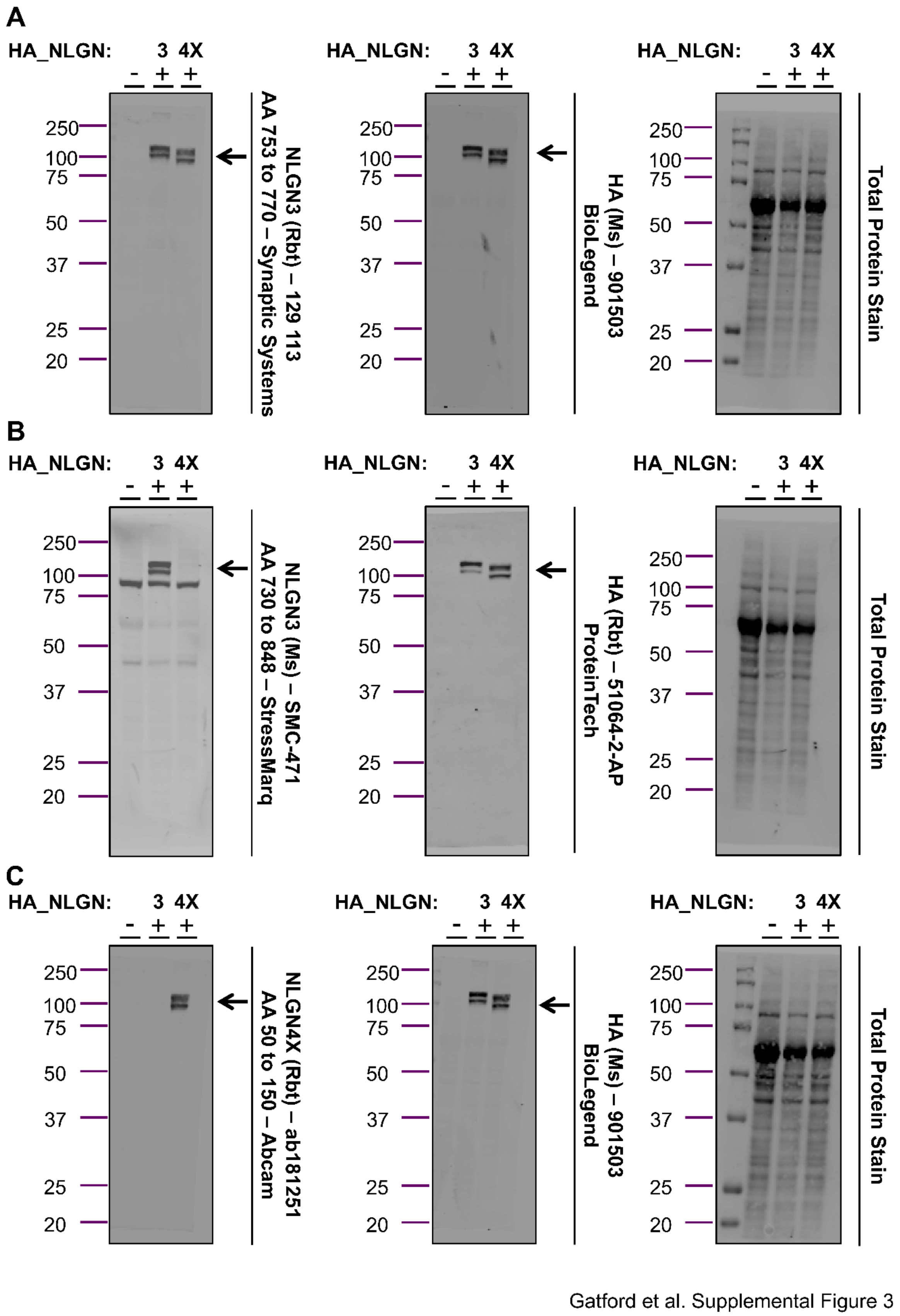


**Supplemental Figure 3** – NLGN3 and NLGN4X antibodies have a degree of cross reactivity. (A) Representative western blots showing cross reactivity for the NLGN3

(Rabbit) antibody to NLGN3 AA 753 to 770 with NLGN4X. (B) Representative western blots showing cross reactivity for the NLGN3 (Mouse) antibody to NLGN3 AA 730 to 848 with NLGN4X. (C) Representative western blots showing cross reactivity for the NLGN4X (Rabbit) antibody to NLGN4X AA 50 to 150 with NLGN3.

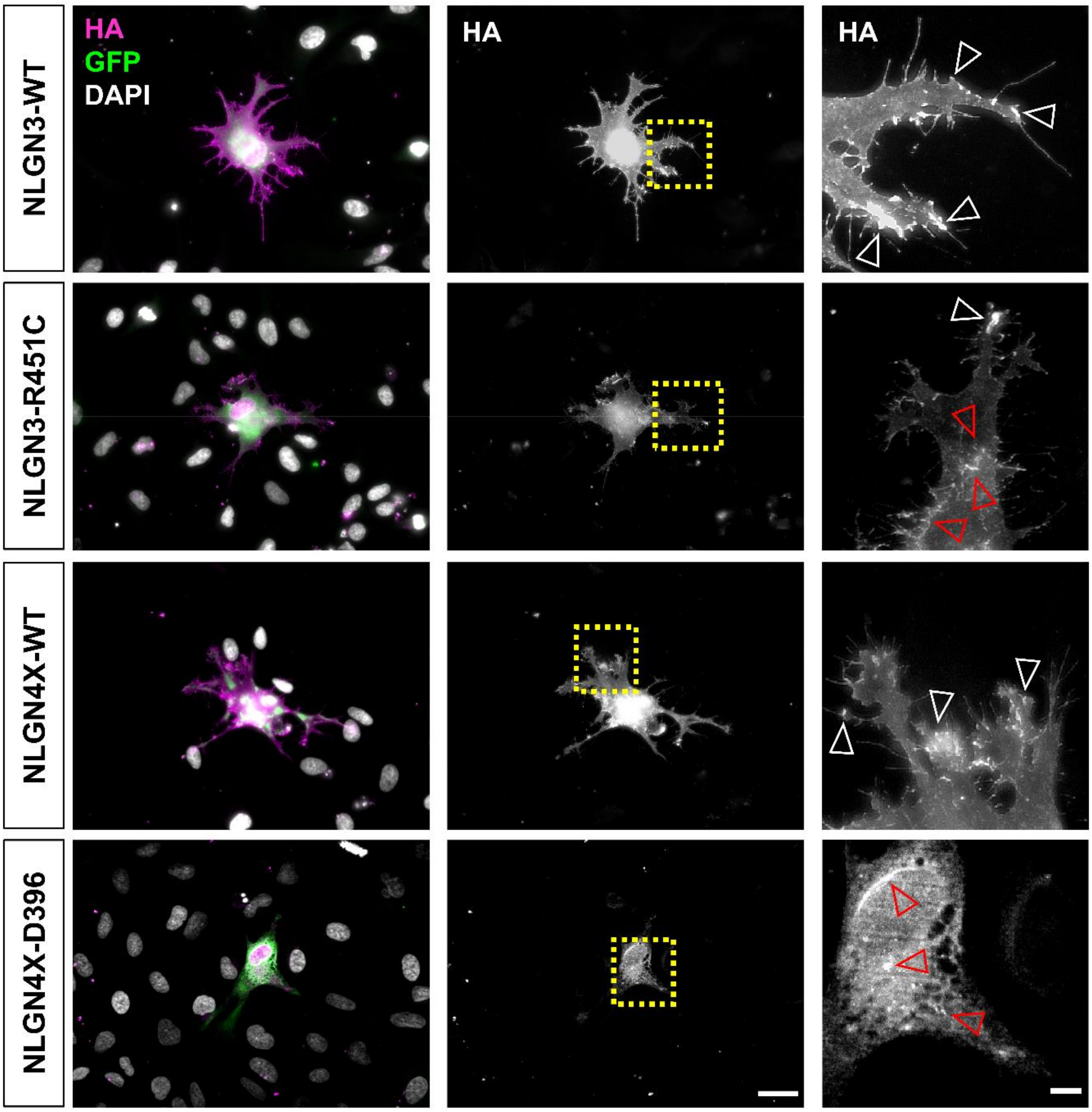


Gatford et al. Supplemental Figure 4

**Supplemental Figure 4** – Representative epifluorescent images of CTX0E18 human neural progenitor cells showing wildtype NLGN3/4X protein localising in growth cones of protrusions (white open arrows) while mutant variant proteins mislocalise to the cytosol (red open arrows). Scale bar = 25µm (full cell), 5µm (magnified).

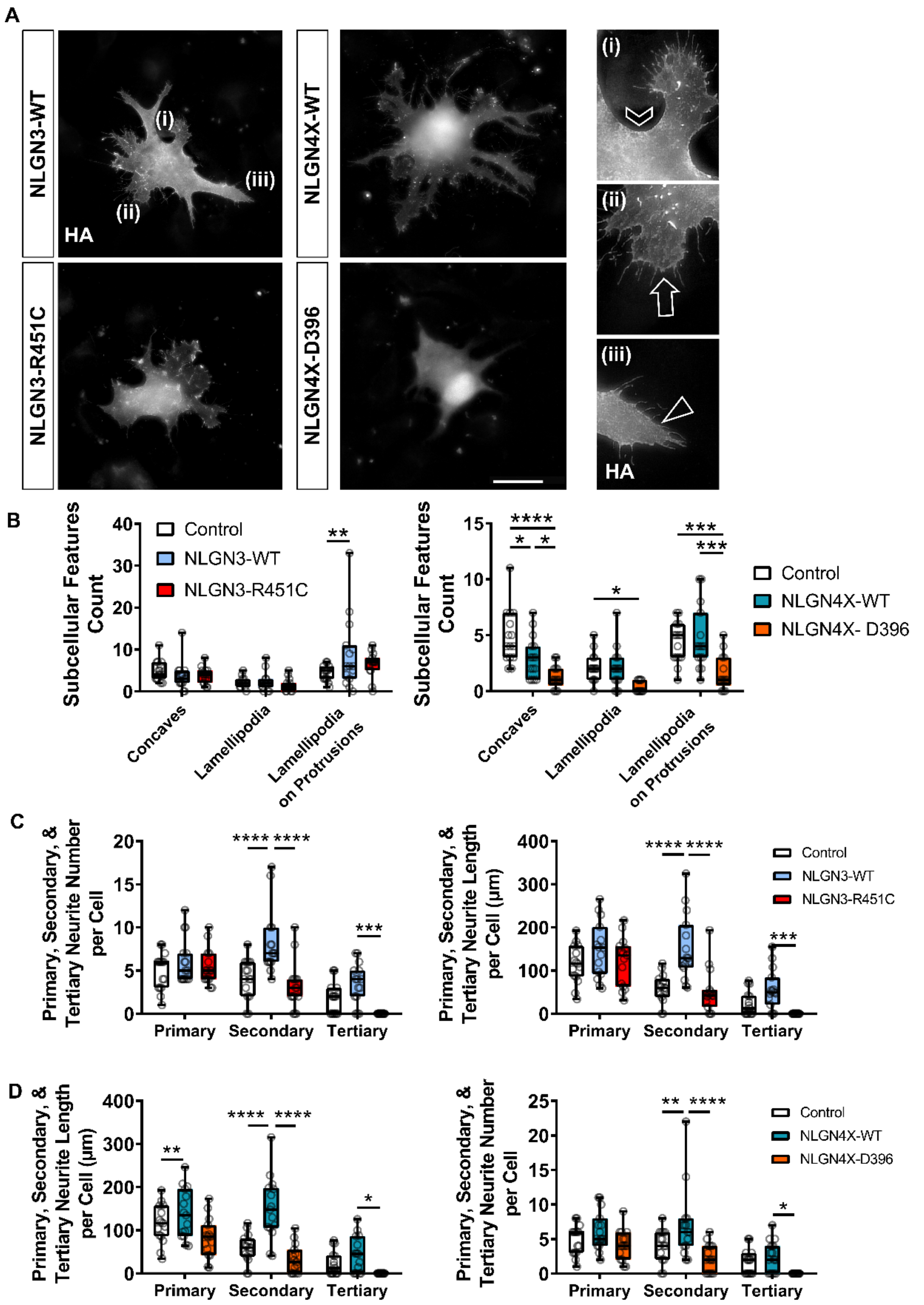


Gatford et al. Supplemental Figure 5

**Supplemental Figure 5** – NLGN3-WT, NLGN4X-WT, and their mutant forms exert differential effects on subcellular features and neurite outgrowth. (A) Representative images showing ectopic NLGN3 and NLGN4X expression induces cell membrane structural changes which are not induced by ectopic expression of their mutant variants. Chevron = concave, arrow = lamellipodia, triangle = lamellipodia on protrusion. Scale bar = 25µm. (B) Data showing ectopic NLGN3 expression significantly increases the number of lamellipodia on protrusions. One-way ANOVA: $F(4,126)=2.58$, $p=0.041$, Bonferroni: $t(126)=3.39$, $p=0.003$, $n=15$. Additionally, ectopic NLGN4X expression significantly decreased the number of concaves, but increased the number of lamellipodia and lamellipodia on protrusions. Concaves: One-way ANOVA: $F(4,120)=23.84$, $p<0.0001$, Bonferroni: $t(120)=5.13$, $p<0.0001$, $n=15$; lamellipodia: Bonferroni: $t(120)=2.52$, $p=0.039$, $n=15$; lamellipodia on protrusions: Bonferroni: $t(120)=3.80$, $p=0.0007$, $n=15$.

A

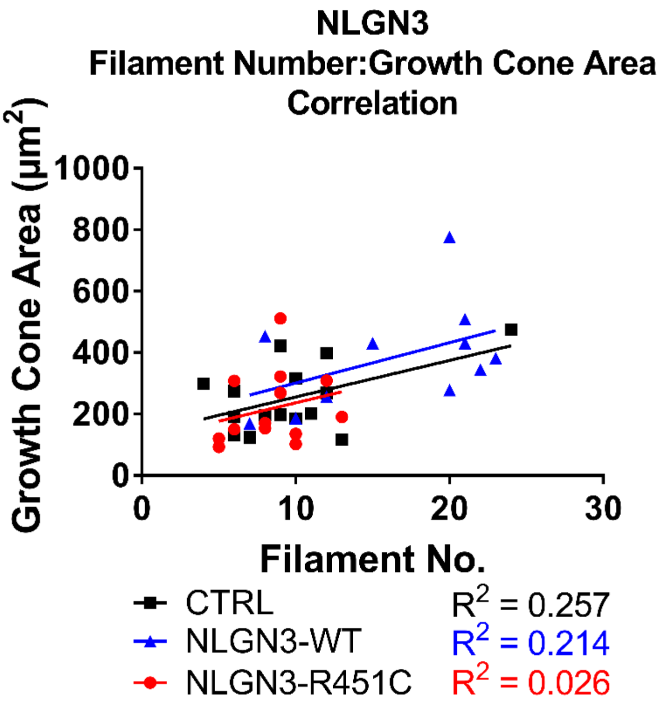


B

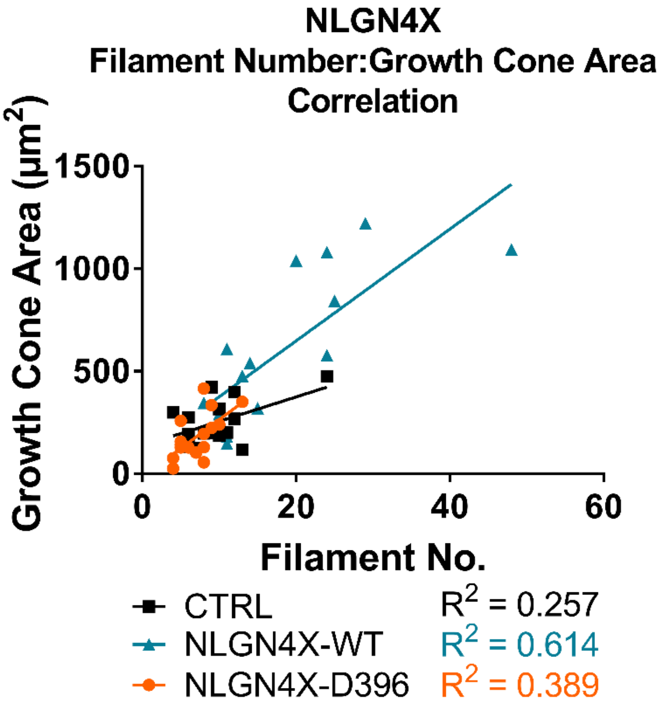


Gatford et al. Supplemental Figure 6

**Supplemental Figure 6** – (A) Correlation data showing filament number and growth cone area are correlated in growth cones ectopically expressing wildtype NLGN3 and NLGN3-R451C. (B) Correlation data showing filament number and growth cone area are correlated in growth cones ectopically expressing wildtype NLGN4X and NLGN4X-D396.

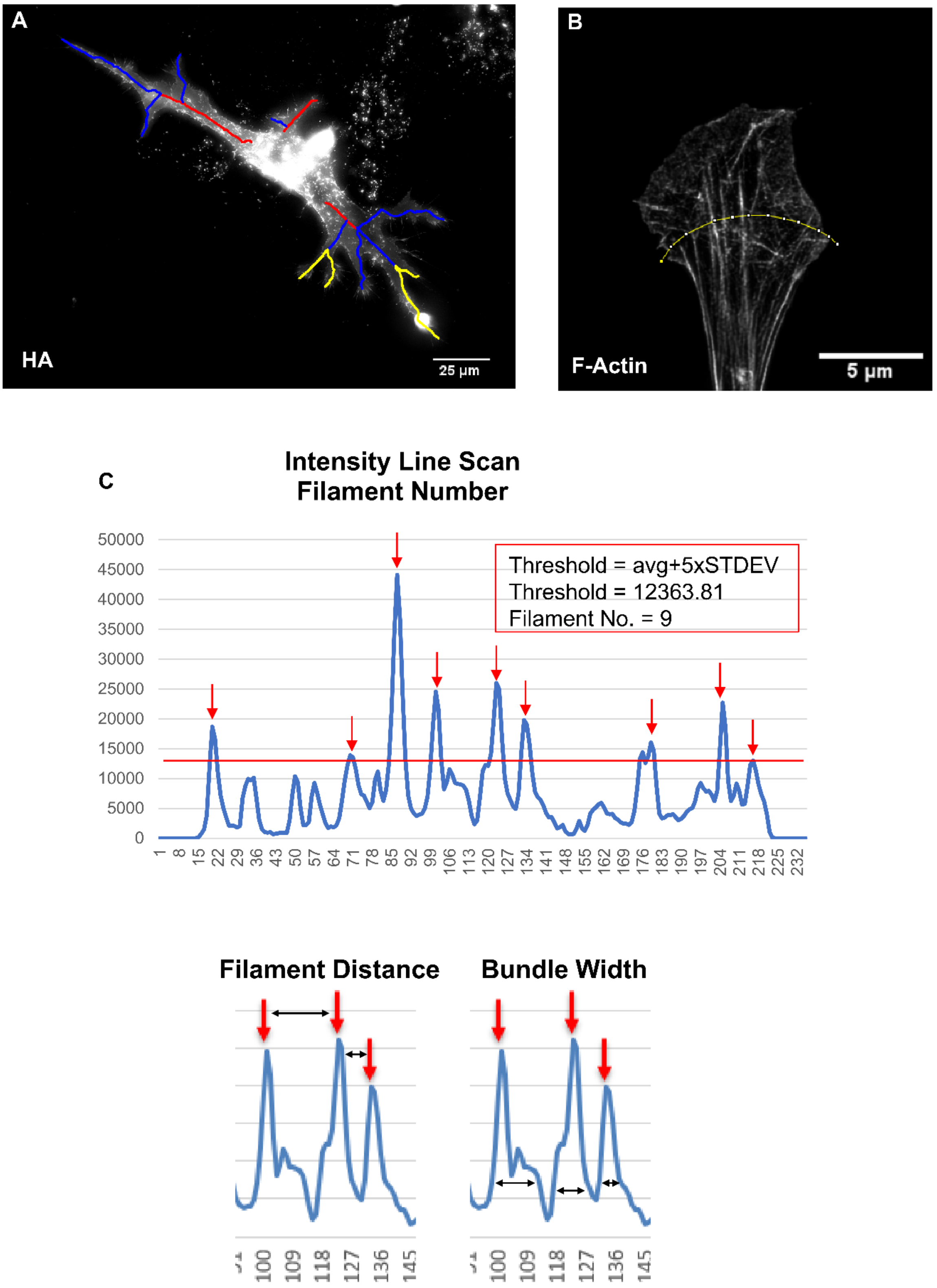


Gatford et al. Supplemental Figure 7

**Supplemental Figure 7** – Example images and diagrams illustrating how neurites and actin filaments were quantified. (A) Example image of a CTX0E16 human neural progenitor cell ectopically expressing wildtype NLGN3 with overlaid neurite tracings; red, primary neurite; blue, secondary neurite; yellow, tertiary neurite. Scale bar = 25µm. (B) Example image of F-actin in a growth cone from an untransfected control CTX0E16 immature neuron with a line scan overlay showing how actin filaments were detected. Scale bar = 5µm (C) Example graphs showing data output from the line scan in (B), illustrating how the background threshold was calculated, how filaments were counted, how filament distance was calculated, and how bundle width was measured.
